# Metabolically Activated Proteostasis Regulators Protect Against Glutamate Toxicity by Activating NRF2

**DOI:** 10.1101/2021.06.29.450364

**Authors:** Jessica D. Rosarda, Kelsey R. Baron, Kayla Nutsch, Michael J. Bollong, R. Luke Wiseman

## Abstract

The extracellular accumulation of glutamate is a pathologic hallmark of numerous neurodegenerative diseases including ischemic stroke and Alzheimer’s disease. At high extracellular concentrations, glutamate causes neuronal damage by promoting oxidative stress which can lead to cellular death. This has led to significant interest in developing pharmacologic approaches to mitigate the oxidative toxicity caused by high levels of glutamate. Here, we show that the small molecule proteostasis regulator AA147 protects against glutamate-induced cell death in a neuronal-derived cell culture model by reducing intracellular levels of reactive oxygen species. While originally developed as an activator of the ATF6 arm of the unfolded protein response, we show AA147-dependent protection against glutamate toxicity is primarily mediated through activation of the NRF2-regulated oxidative stress response. We demonstrate that AA147 activates NRF2 through a mechanism involving metabolic activation to a reactive electrophile and covalent modification of KEAP1 – a mechanism analogous to that involved in AA147-dependent activation of ATF6. These results define the potential for AA147 to protect against glutamate induced oxidative toxicity and highlight the potential for metabolically-activated proteostasis regulators like AA147 to activate both protective ATF6 and NRF2 stress-responsive signaling pathways to mitigate oxidative damage associated with diverse neurologic diseases.

## INTRODUCTION

Glutamate is an essential excitatory neurotransmitter involved in nervous system function. The controlled release of glutamate into the synapse is critical for neuronal signaling (1, 2). However, acute or chronic events which cause pathologic depolarization of neuronal cell membranes lead to an uncontrolled release of glutamate into the extracellular space, causing aberrant excitotoxic and oxidative signaling that can lead to cell death (1-3). High levels of extracellular glutamate trigger a cascade of excitatory signaling through the excessive stimulation of neuronal NMDA receptors which release additional glutamate into the synapse (2, 4). Neurons reduce extracellular glutamate levels by reversing the function of the XC-antiporter, which begins to import glutamate and export cystine (5, 6). Cystine is a vital precursor to the intracellular antioxidant glutathione and prolonged reversal of the XC-system depletes glutathione stores (7). Therefore, high levels of extracellular glutamate result in decreased antioxidant capacity within the cell, causing oxidative stress that can lead to cell death independent of NMDA receptor activation (6, 8). Limiting excitoxicity using NMDA receptor antagonists protects against neurodegeneration in multiple neurologic disorders, including human and mouse models of Alzheimer’s disease and ischemic stroke (3, 9, 10). However, inhibition of these receptors can be problematic, as it also disrupts physiologic excitatory signaling (3, 11). An alternative approach to treat these disorders is to limit the cell death caused by glutamate-induced oxidative damage (12, 13).

We recently identified the compound AA147, which is protective against ROS mediated damage caused by ischemia and reperfusion (I/R) injury in both cells and mice (14, 15). Administration of AA147 improved outcomes in mouse models of cardiac and kidney I/R injury. Further, AA147 administered either prior to the onset of ischemia or at the time of reperfusion reduced both the infarct size and neurological dysfunction in mice subjected to cerebral I/R (14). Glutamate toxicity is a major contributor to neurologic damage following an ischemic stroke, which suggested that AA147 could decrease cerebral I/R damage by ameliorating glutamate toxicity (3, 11).

AA147 was originally developed as a pharmacologic activator of the Activating Transcription Factor 6 (ATF6) signaling pathway within the unfolded protein response (UPR) (15, 16). Activation of ATF6 upregulates a transcriptional response during conditions of endoplasmic reticulum (ER) stress through a process involving increased trafficking of full-length ATF6 to the Golgi and subsequent proteolytic release of the active N-terminal ATF6 transcription factor domain by site 1 (S1P) and site 2 (S2P) proteases (16, 17). Upon nuclear localization, ATF6 induces expression of multiple ER proteostasis factors including protein chaperones, such as BiP, GRP94, and PDIA4, as well as redox factors, such as HMOX1 (15, 16, 18, 19). AA147 induces the nuclear translocation of ATF6 through a mechanism involving compound metabolic activation to a reactive electrophile and subsequent covalent modification of a subset of ER-localized protein disulfide isomerases (PDIs) involved in regulating the trafficking of ATF6 to the Golgi (20). This mechanism allows AA147 to preferentially activate the ATF6 arm of the UPR both in cell culture models and in vivo (15, 16).

ATF6 transcriptional signaling is protective in models of etiologically diverse diseases, making this pathway an attractive therapeutic target for disease intervention (14, 16, 21-24). Consistent with this, pharmacologic activation of ATF6 with AA147 is protective in models of numerous diseases. For example, AA147-dependent ATF6 activation is protective in mouse models of myocardial infarction and cardiac arrest, as well as iPSC-derived models of the eye disease achromatopsia (14, 24, 25). However, AA147 can also promote protection through ATF6-independent mechanisms. AA147-dependent covalent modification of PDIs, an upstream step involved in AA147-dependent ATF6 activation (20), is sufficient to reduce secretion and toxic aggregation of amyloidogenic immunoglobulin light chains associated with Light Chain Amyloidosis independent of ATF6 signaling (26). Further, AA147 protects the liver against viral infection through an ATF6-independent mechanism (27). These results highlight that apart from ATF6 activation, AA147 can also protect against diverse pathologic insults through mechanisms independent of ATF6 activity.

Here, we sought to define the potential for AA147 to protect against glutamate-induced oxidative toxicity in HT22 cells, an immortalized cell line derived from hippocampal neurons that lack the NMDA receptors required to drive the excitatory release of glutamate to the extracellular space (8). Treatment of HT22 cells with glutamate induces oxidative stress independent of excitatory signaling, making this a widely used model to develop pharmacologic approaches to mitigate the oxidative toxicity caused by glutamate (5, 8, 28). We show that AA147 protects HT22 cells against glutamate-induced oxidative toxicity. Intriguingly, this protection is only partially dependent on ATF6 activation. Instead, AA147-dependent protection against glutamate-induced oxidative toxicity is primarily mediated through compound-dependent activation of the NRF2 oxidative stress response. Interestingly, structure-activity relationships indicate that AA147 activates NRF2 through a mechanism involving compound metabolic activation and covalent targeting of the NRF2 regulatory protein KEAP1, a mechanism analogous to that involved in AA147-dependent ATF6 activation (20). These results demonstrate that AA147 offers a unique opportunity to activate both adaptive ATF6 and NRF2 transcriptional signaling, revealing further insights into the molecular basis for protection afforded by this compound. Further, our results demonstrate the broad potential for AA147 and related compounds to mitigate oxidative damage induced by pathologic insults through the coordinated regulation of two protective stress-responsive signaling pathways.

## RESULTS & DISCUSSION

### AA147 protects against glutamate-induced oxidative toxicity

We sought to define the potential for AA147 to reduce glutamate induced oxidative toxicity in HT22 cells. We initially confirmed that AA147 activated the ATF6-selective ERSE-luciferase reporter (ERSE-LUC)(15) in HT22 cells with an EC50 of 3.6 µM (**Fig. S1A**). Further, we showed that AA147 did not significantly influence HT22 cell viability (**Fig. S1B**). These results are consistent with AA147 activity observed in other cell models (15) and demonstrates that AA147 is active in HT22 cells. Next, we assessed whether AA147 reduces glutamate-induced oxidative toxicity in HT22 cells. Initially, we used the MTT assay to monitor the viability of HT22 cells pre-treated with AA147 and then challenged with glutamate for 24 h (**Fig. 1A**). Addition of AA147 concurrently with the glutamate challenge did not improve viability of glutamate-treated cells (**Fig. S1C**). However, pre-treatment with AA147 for 6 or 16 h prior to the glutamate challenge increased viability of glutamate-treated HT22 cells by 2- or 3-fold, respectively (**Fig. 1B, C**). Similar results were observed when monitoring glutamate-induced cell death in HT22 cells by Annexin V and propidium iodide (PI) staining, where pretreatment with AA147 for 6 or 16 h reduced the population of Annexin V/PI positive cells (**Fig. 1D,E, Fig. S1D,E**). Again, 16 h pre-treatment proved to be most effective at mitigating cell death.

**Figure 1.**
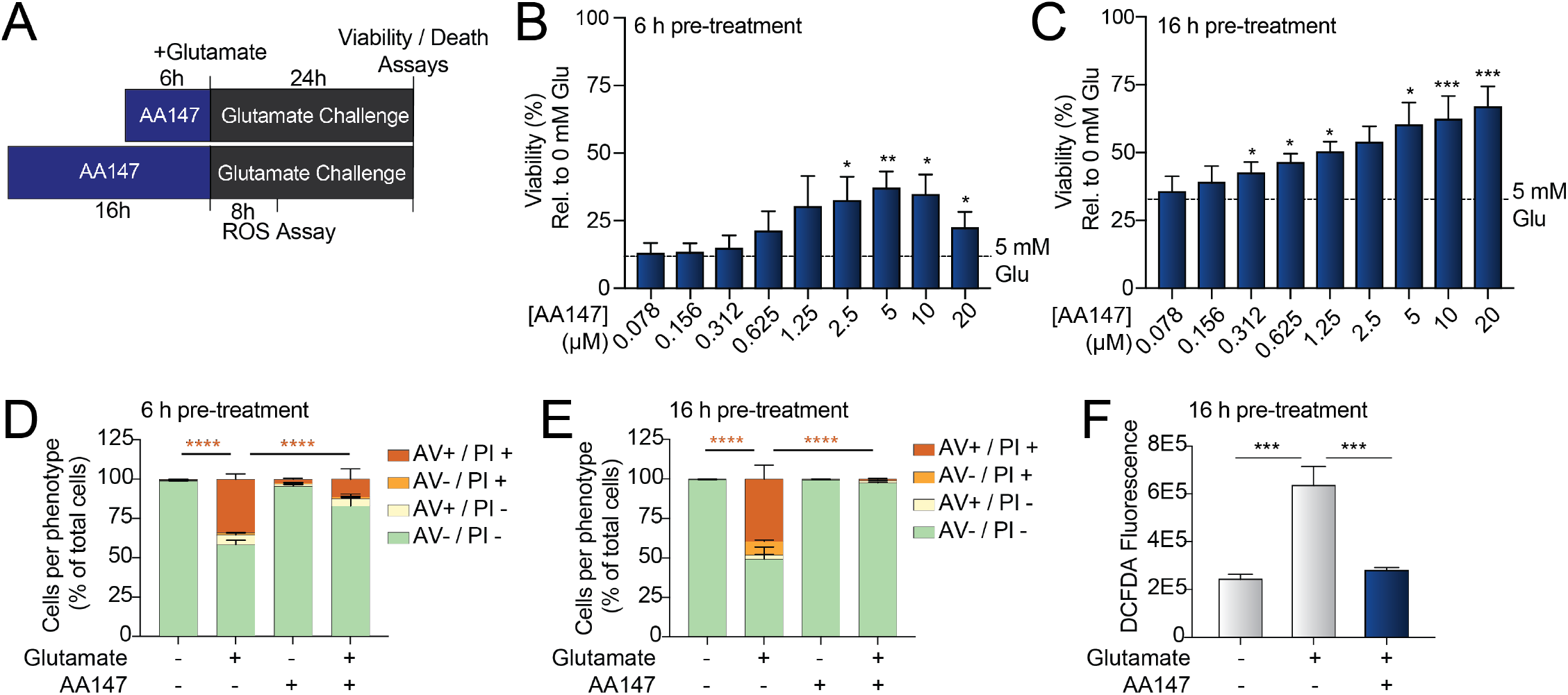
AA147 protects against glutamate-induced oxidative toxicity in HT22 cells. **A**. Schematic of the AA147 pre-treatment conditions and glutamate challenge. **B,C**. Viability, measured by MTT assay, of HT22 cells pre-treated with the indicated dose of AA147 for 6 h (**B**) or 16 h (**C**) and then challenged with glutamate (5 mM) for 24 h. Viability is shown as a percent relative to vehicle treated cells where glutamate was not added. Error bars show SEM for n=4 (**B**) or n=5 (**C**) replicates. *p<0.05, **p<0.01, ***p<0.005 for two-tailed paired Student’s t-test. **D,E**. Quantification of the percent of HT22 cells pre-treated with AA147 for 6 or 16 h and then challenged with glutamate (5 mM) for 24 h stained with Annexin V (AV) and/or propidium iodide (PI). Error bars show SEM for n=3 replicates. ****p<0.0001 for ordinary one-way ANOVA with Tukey correction for multiple comparisons. **F**. Geometric mean of CM-H2DCFDA fluorescence of HT22 cells pre-treated with AA147 (10 µM) for 16 h and then challenged with glutamate (5 mM) for 8 h, as indicated. Error bars show SEM for n=3 replicates. ***p<0.005 for ordinary one-way ANOVA with Tukey correction for multiple comparisons.

AA147 reduces toxicity induced by oxidative stress in several cell types by decreasing reactive oxygen species (ROS)-associated damage (14). Thus, we sought to determine if AA147 reduced ROS levels in glutamate-treated HT22 cells. HT22 cells show a significant increase in ROS 8 h after the addition of glutamate, as measured by DCFDA fluorescence (**Fig. 1F**), consistent with published results (28). Pretreatment with AA147 for 16 h significantly reduced DCFDA fluorescence in glutamate challenged cells, indicating that AA147 reduces ROS accumulation in these cells. Collectively, these results demonstrate that AA147 attenuates glutamate-induced oxidative toxicity in HT22 cells.

### The 2-p-amino cresol substructure of AA147 is required for protection against glutamate toxicity

AA147 consists of a 2-*p*-amino cresol moiety (designated as the A-Ring) linked to an aromatic B-ring via a hydrocarbon linker (**Fig 2A**) (15, 20). Previous studies demonstrated that the 2-*p*-amino cresol moiety is metabolically activated by ER oxidases to form reactive electrophiles such as a quinone methide (AA147-QM) or quinone imine (AA147-QI)(**Fig. 2B**)(20). These electrophilic forms of AA147 covalently modify reactive cysteines on proteins predominantly localized to the endoplasmic reticulum (ER) (20). Initially, we asked whether AA147 covalently modified proteins in HT22 cells. We used an analog of AA147 with an alkyne handle on the B-ring (AA147^alk^)(**Fig. S2A**), which allows monitoring of covalently modified proteins by ‘click chemistry’ (20). AA147^alk^ protected against glutamate-induced toxicity in HT22 cells (**Fig. S2B**). We showed that AA147^alk^ covalently modified proteins in HT22 cells by conjugating a rhodamine-azide to AA147^alk^-modified proteins using a copper-catalyzed alkyne-azide cycloaddition reaction (**Fig. S2C**). Co-treatment with excess AA147 blocked AA147^alk^ labeling, indicating that these two compounds modify the same proteins in HT22 cells. These results demonstrate that AA147 covalently modifies proteins in HT22 cells, mirroring results observed in other cell types (20, 26).

**Figure 2.**
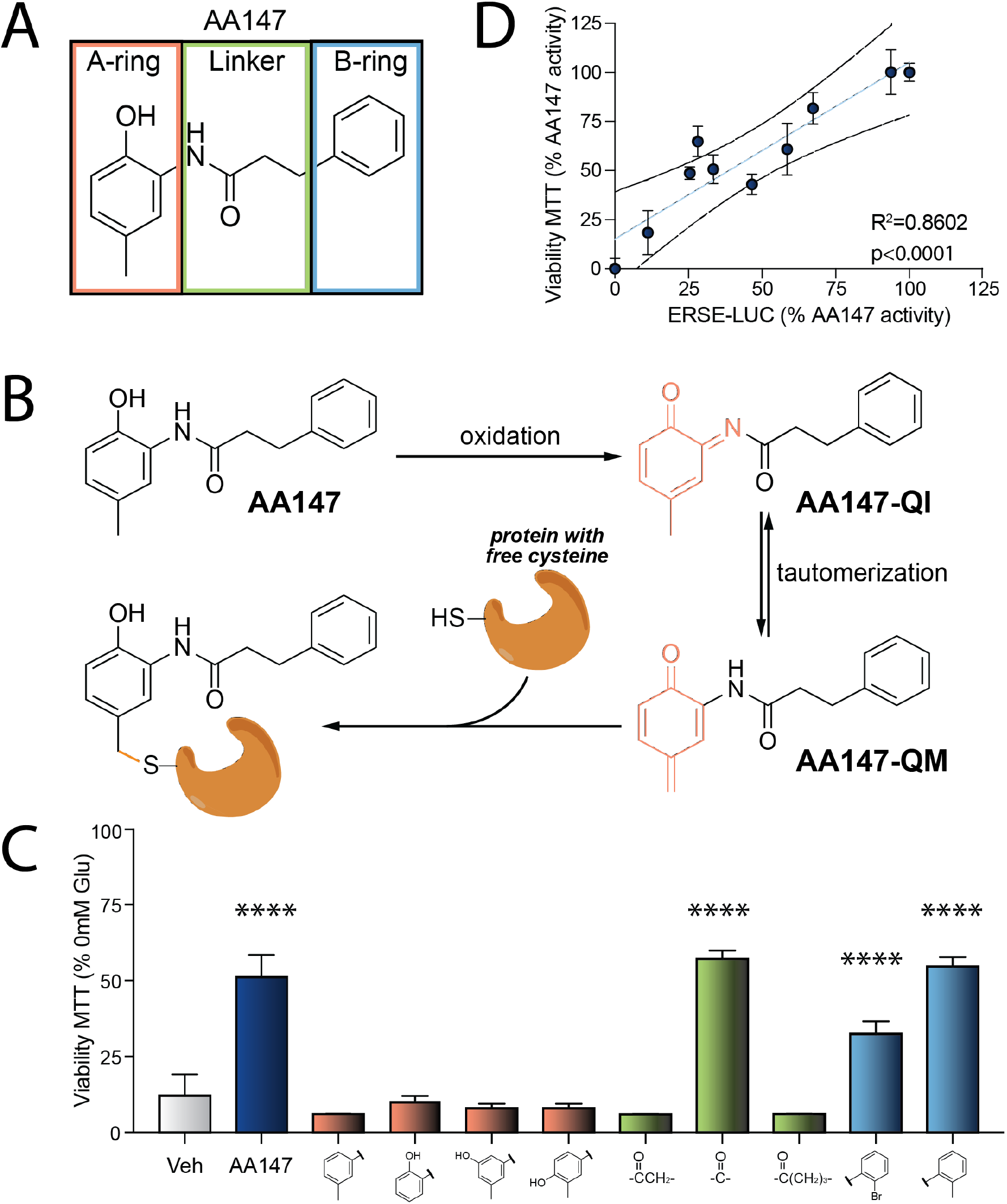
The 2-*p*-amino cresol substructure of AA147 is required to protect HT22 cells against glutamate induced oxidative toxicity. **A**. Structure of AA147 with its three components – the A-ring, linker, and B-ring, highlighted. The A-ring (orange) contains the 2-*p*-amino cresol moiety. **B**. Schematic showing the mechanism of AA147 metabolic activation to a quinone-imine (AA147-QI) or quinone methide (AA147-QM) and subsequent covalent protein modification. Panel adapted from (20). **C**. Viability, measured by MTT, of HT22 cells pre-treated with the indicated AA147 analog (10 µM) for 16 h and challenged with glutamate (5 mM) for 24 h. Viability is shown as a percent of HT22 cells pre-treated with respective treatments in the absence of glutamate. Error bars show SD for n=3 replicates ****p<0.0001 for ordinary one-way ANOVA against vehicle treated cells with Dunnett correction for multiple comparisons. **D**. Correlation between viability of HT22 cells pre-treated with respective analogs for 6h and then challenged by glutamate for 24 h (measured by MTT) and activation of the ATF6-selective ERSE-LUC reporter in transiently transfected HT22 cells. R^2^ and p-value calculated from a simple linear regression.

Next, we tested the dependence of AA147-dependent protection against glutamate-induced oxidative toxicity on the metabolic activation of the A-ring using a series of AA147 analogs. Compounds with disruptions in the 2-*p*-amino cresol A-ring did not protect against glutamate-induced toxicity (**Fig. 2C**). In contrast, the B-ring of AA147 was more tolerant to modification and many of these analogs showed effective protection. This is an analogous structure-activity relationship to that previously shown to regulate ATF6 activation by AA147 in HEK293 cells (20). Consistent with this, the protection afforded by a 6 h pre-treatment with AA147 analogs linearly correlates with their activation of the ATF6-selective ERSE-LUC reporter in HT22 cells (**Fig. 2D**). These results indicate that AA147 protects against glutamate-induced toxicity in HT22 cells through a mechanism similar to that required for ATF6 activation (**Fig. 2B**)(20, 26).

### AA147-dependent ATF6 activation only modestly contributes to protection against glutamate-induced toxicity observed in HT22 cells

AA147 protects cardiomyocytes from oxidative insults through activation of ATF6 (14). Thus, we sought to define the dependence of AA147-dependent protection against glutamate-induced oxidative toxicity on ATF6 activation. Initially, we used an inhibitor of ATF6 activation, the S1P inhibitor PF429242 (S1Pi)(29), to define the importance of AA147-dependent ATF6 activation on the protection observed in glutamate-treated HT22 cells. We confirmed that S1Pi inhibited AA147-dependent induction of the ATF6 target *BiP/Hspa5* in HT22 cells by qPCR and immunoblotting (**Fig. S3A** and **Fig. S3B**). Next, we monitored the viability of HT22 cells pretreated with AA147 and S1Pi for 6 h or 16 h and then challenged with glutamate. Co-treatment with S1Pi modestly attenuated the AA147-dependent protection observed following a 6 h pretreatment (**Fig. 3A**). Similarly, co-treatment with S1Pi moderately impacted AA147-dependent reductions in ROS (**Fig. S3C**). These results suggested that AA147-dependent ATF6 activation contributes to the protection observed following 6 h pretreatment. However, co-treatment with S1Pi did not affect AA147-dependent improvements in viability observed following a 16 h pretreatment (**Fig. 3B**). shRNA depletion of *Atf6* also did not reduce protection against glutamate toxicity in HT22 cells pre-treated for 16 h with AA147 (**Fig. 3C** and **Fig. S3D**). This indicates that AA147-dependent ATF6 activation does not contribute to the protection against glutamate-induced toxicity at this timepoint. Instead, AA147 must promote protection through an alternative mechanism.

**Figure 3.**
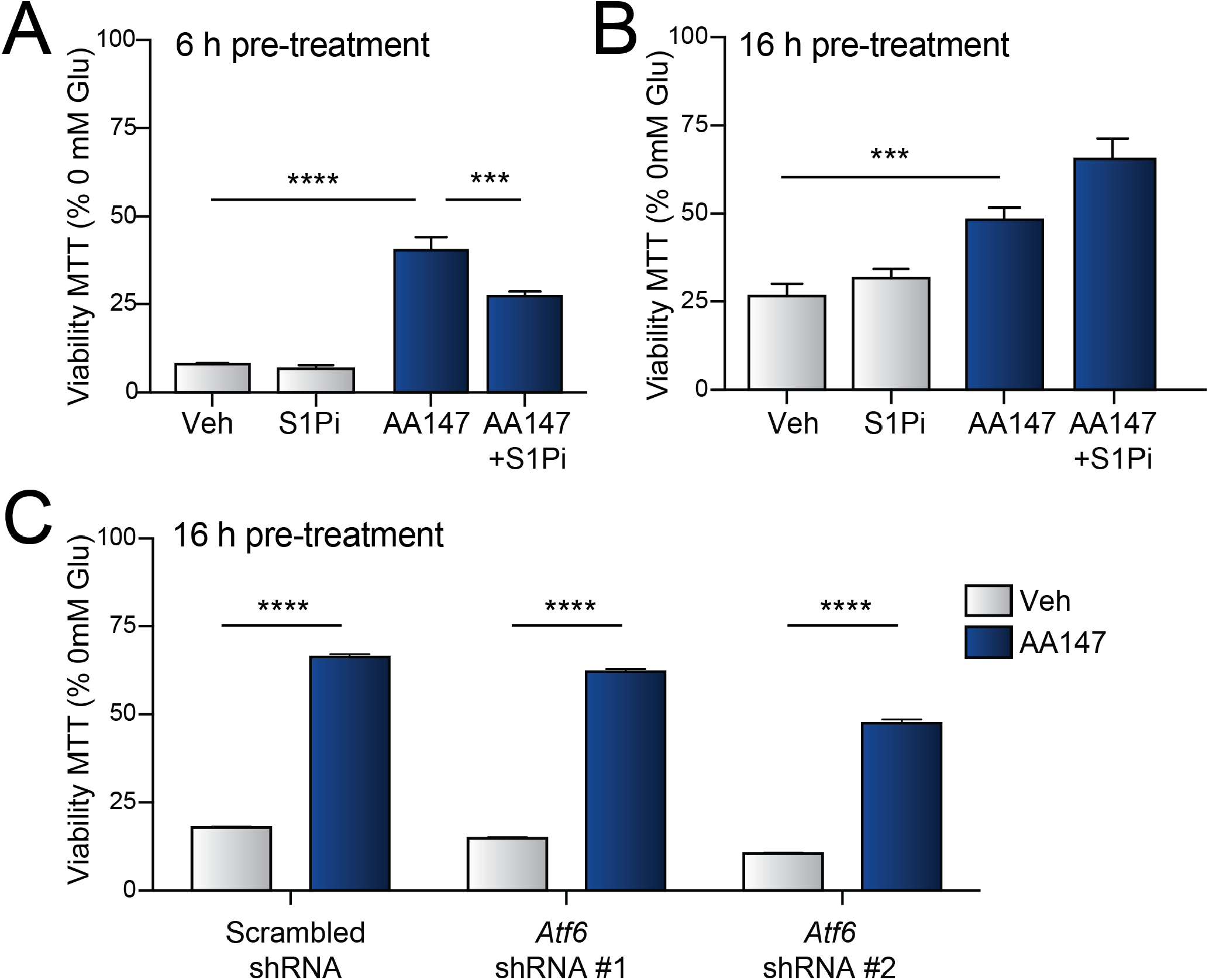
AA147-dependent activation of ATF6 modestly contributes to the AA147-dependent protection of HT22 cells against glutamate induced oxidative toxicity. **A,B**. Viability, measured by MTT, of HT22 cells pre-treated with AA147 (10 µM) for 6 h (**A**) or 16 h (**B**) in the presence or absence of S1Pi (10 µM) and then challenged with glutamate (5 mM) for 24 h. Viability is shown as percent relative to cells treated with the respective treatment in the absence of glutamate. Error bars show SD for n=3. ***p<0.001, ****p<0.0001 for two-way ANOVA with Tukey correction for multiple testing. **C**. Viability, measured by MTT assay, of HT22 cells expressing scrambled or *Atf6* shRNA pre-treated for 16 h with AA147 (10 µM) and then challenged with glutamate (5 mM) for 24 h. Viability is shown as percent relative to cells treated with the respective treatment in the absence of glutamate. Error bars show SD for n=3. ****p<0.0001 for two-way ANOVA with Tukey correction for multiple testing.

### AA147 activates the oxidative stress response in HT22 cells

To better define the mechanistic basis for AA147-dependent protection against glutamate induced oxidative toxicity, we performed RNA sequencing (RNAseq) on HT22 cells treated with vehicle or AA147 for 16 h (**Table S1**). Despite observing a robust induction of the ATF6 target gene *BiP* following 6 h treatment (**Fig. S3A**), RNA-seq showed that *BiP* expression was not increased following 16 h pretreatment with AA147 in these cells (**Table S1**). We confirmed this result by qPCR (**Fig. S4A**). This is consistent with the transient AA147-dependent activation of ATF6 signaling observed in other cells (15). However, we observed significant increases in the expression of numerous oxidative stress responsive genes including prolactins (e.g., *Prl2c2, Prl2c3*) and glutathione transferases (e.g., *Gsta1, Gsta4*) in AA147-treated HT22 cells (**Fig. 4A**)(30-32). GO analysis similarly showed increases in oxidative stress-associated biological pathways including glutathione transferase activity and prolactin receptor binding (**Fig. S4B**). Further, when monitoring the expression of established gene sets comprised of transcriptional targets of different stress-responsive signaling pathways (33), only oxidative stress response target genes showed a coordinated upregulation, as compared to control genes, in AA147-treated HT22 cells (**Fig. 4B, Table S2**). Notably, transcriptional targets of ATF6 or other arms of the UPR (i.e., XBP1s and PERK) were not induced in HT22 cells treated for 16 h with AA147 (**Fig. 4B, Table S2**), again reflecting the transient nature of AA147-dependent activation of ATF6 (15). These results suggest that AA147 induces an oxidative stress response in HT22 cells.

**Figure 4.**
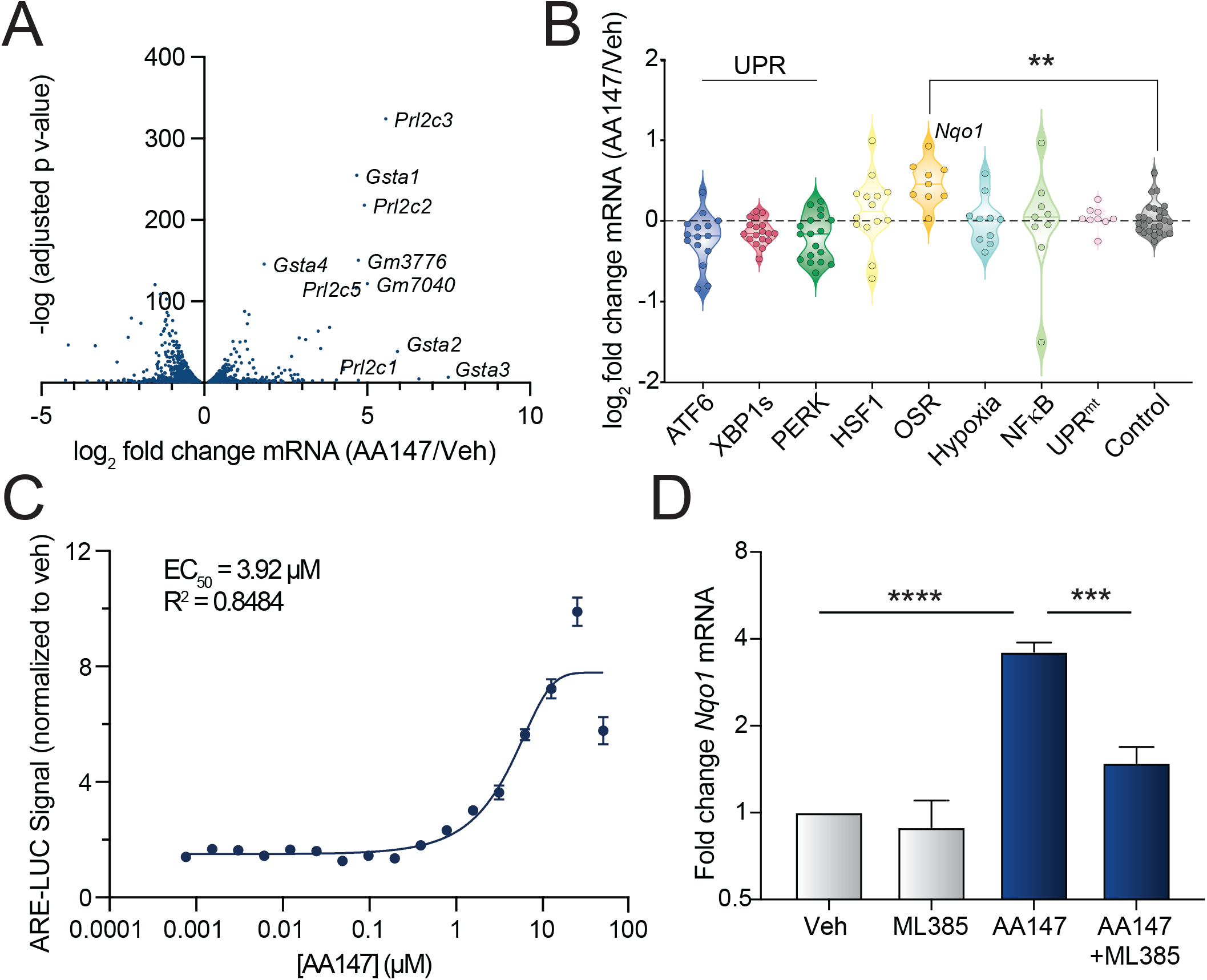
AA147 induces NRF2-dependent upregulation of oxidative stress response genes in HT22 cells. **A**. Plot showing -log adj p-value vs log2 fold change (AA147/Veh) for genes identified in RNA-seq analysis of HT22 cells treated with vehicle or AA147 (10 µM) for 16 h. Select glutathione transferase and prolactin genes are indicated. Complete RNAseq data is shown in **Table S1. B**. Graph showing expression of sets of genes regulated downstream of the ATF6, XBP1s, or PERK arms of the UPR, the HSF1-regulated heat shock response, the oxidative stress response (OSR), the hypoxia stress response, NFκB inflammatory signaling, and the mitochondrial unfolded protein response (UPR^mt^). A subset of control genes is also shown. Gene sets are defined as previously described (33) and are shown in **Table S2**. **p<0.01 for one-way ANOVA comparing expression to control genes. **C**. Luminescence in HT22 cells transiently expressing the ARE-LUC reporter and treated with AA147 (10 µM) for 16 h. Luminescence is shown as fold change relative to vehicle. Error bars show SEM for n=20 replicates across two independent experiments. **D**. Expression of the NRF2 target gene *Nqo1*, measured by qPCR, in HT22 cells treated with vehicle or AA147 (10 µM) in the presence or absence of ML385 (5 µM) for 16 h. Error bars show SEM for n=3. ***p<0.001, ****p<0.0001 for two-way ANOVA with Tukey correction for multiple testing.

The oxidative stress response is primarily regulated by the transcription factor NRF2, which binds to antioxidant response elements (ARE) sequences within the promoter region of target genes to induce their expression (34, 35). NRF2 activity protects against multiple different types of oxidative insults, including glutamate-induced toxicity (36, 37). Thus, we sought to determine whether AA147 was activating NRF2 in HT22 cells. Initially, we showed that AA147 increased expression of an NRF2-selective ARE-LUC reporter in HT22 cells (**Fig 4C**)(34). AA147 activated this ARE-LUC reporter with an EC50 of 3.9 µM, which is nearly identical to that observed for activation of the ATF6-selective ERSE-LUC reporter (**Fig. S1A**). Next, we used qPCR to confirm that AA147 induced expression of NRF2 target genes including *Nqo1* and *Gsta4* in HT22 cells (**Fig. 4D** and **Fig. S4C**). Co-treatment with the NRF2 inhibitor compound ML385, which inhibits NRF2 binding to DNA (38), reduced AA147-dependent *Nqo1* and *Gsta4* induction (**Fig. 4C, S4C**). Similar results were observed by immunoblotting (**Fig. S4D**). Further, shRNA-depletion of *Nrf2* blocked AA147-dependent induction of *Gsta4*, but not *BiP*, in HT22 cells (**Fig. S4E**). This indicates that AA147 activates NRF2 signaling in HT22 cells. Previous RNA-seq profiling did not show increased expression of *Nqo1* or other canonical NRF2 target genes in non-neuronal cell lines including HEK293 cells (15). However, AA147 induced a robust increase in NRF2 target genes such as *Nqo1*, as well as the ATF6 target gene *BiP*, in mouse primary cortical neurons (**Fig. S4F**). Further, AA147 upregulated both the NRF2-selective ARE-LUC reporter (**Fig. S4G**) and the ATF6-selective ERSE-LUC reporter in human IMR32 neuroblastoma cells (**Fig. S4H**). In combination with the existing literature (14, 15, 24, 26, 27), these results suggest that AA147 induces transient ATF6 activity broadly across cell types, whereas AA147-dependent NRF2 activation may only be observed in select cell types including the neuronal-derived HT22 and IMR32 cells.

### AA147 covalently modifies KEAP1 to promote NRF2 activation in HT22 cells

NRF2 activity is primarily regulated through its interaction with the redox sensor E3 ligase protein KEAP1 (**Fig. 5A**)(38, 39). In the absence of oxidative stress, KEAP1 promotes the ubiquitination of NRF2 leading to its inactivation by degradation (39). In response to oxidative stress, sensor cysteine residues on KEAP1, such as Cys151, are covalently modified by electrophiles (e.g., ROS), reducing KEAP1-dependent ubiquitination of NRF2. The resulting decrease in NRF2 ubiquitination stabilizes this transcription factor and allows NRF2 to localize to the nucleus and promote transcriptional activity (39, 40). Since AA147 can be metabolically activated to a reactive electrophile that can covalently modify proteins, we predicted that AA147 activates NRF2 through a mechanism involving metabolic activation and covalent modification of KEAP1.

**Figure 5.**
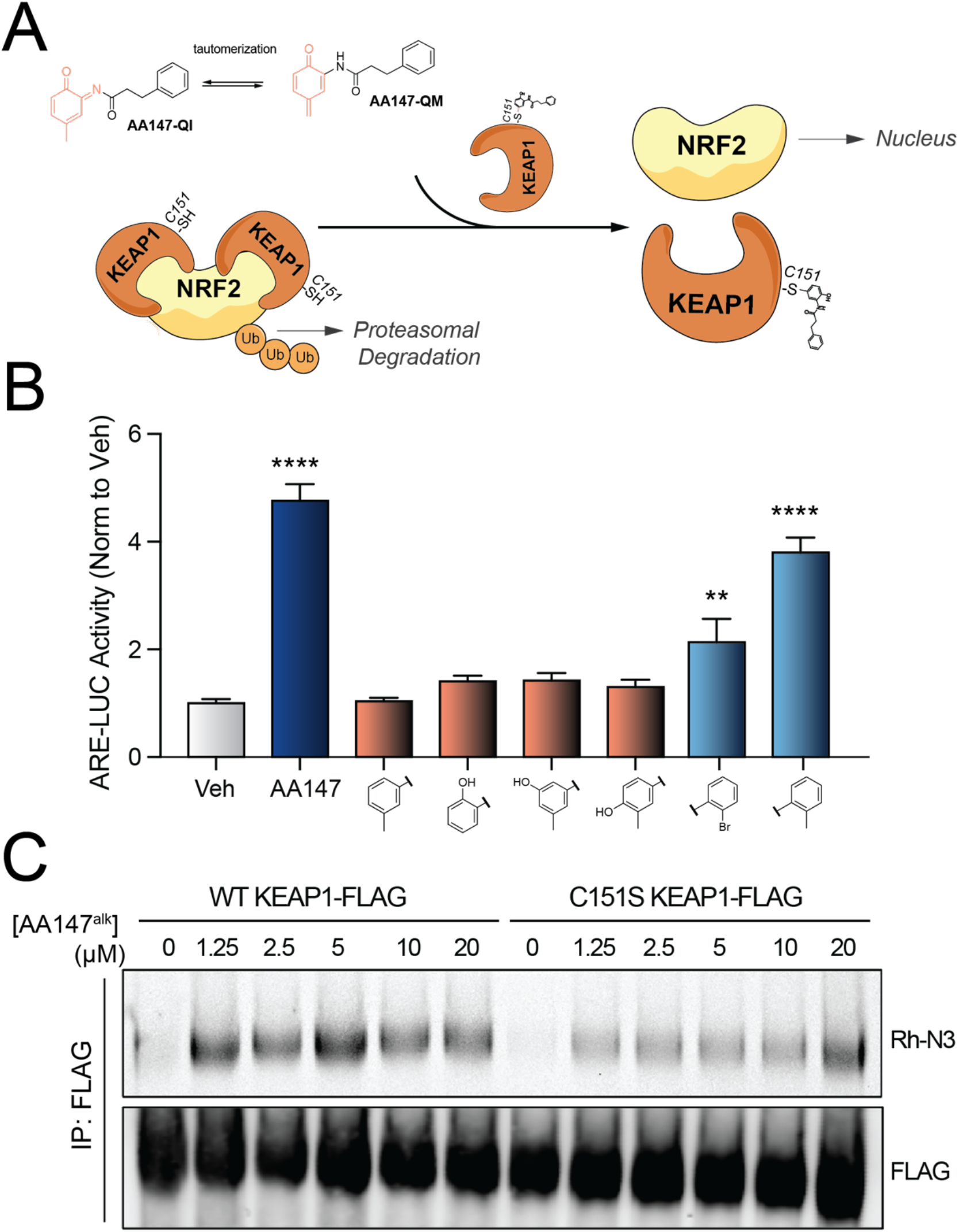
AA147 covalently modifies KEAP1, a regulator of NRF2 transcriptional activity. **A**. Proposed model whereby metabolically activated AA147 covalently modifies Cys151 on KEAP1 to reduce ubiquitination and allow nuclear localization of NRF2 to promote transcriptional activity. **B**. Luminescence in HT22 cells transiently expressing the ARE-LUC NRF2 reporter treated with the indicated AA147 analog (10 µM) for 16 h. Error bars show SEM for n=20 replicates across 2 independent experiments. **p<0.01, ***p<0.001, ****p<0.0001 for ordinary one-way ANOVA against vehicle control with Dunnett correction for multiple comparisons. **C**. Fluorescence image (top) and immunoblot (bottom) of FLAG immunopurifications prepared from HEK293T cells transiently overexpressing wild-type (WT) KEAP^FT^ or C151S KEAP^FT^ and treated for 16 h with the indicated concentration of AA147^alk^. AA147^alk^ modified proteins were conjugated to rhodamine-azide (Rh-N3) by click chemistry.

To test this, we initially monitored the activation of the NRF2 selective ARE-luciferase reporter (ARE-LUC) in HT22 cells treated with AA147 analogs that disrupt compound metabolic activation (20, 34). As expected, analogs which lack the 2-*p*-amino cresol moiety in the AA147 A-ring showed no activation of ARE-LUC (**Fig. 5B**). However, B-ring analogs largely retained ARE-LUC activity. This is a similar structure activity relationship to that observed for activation of the ATF6-selective ERSE-LUC reporter (**Fig. S5A**) and indicates that activation of NRF2 proceeds through the same metabolic activation of the AA147 A-ring required for ATF6 activation (**Fig. 2B**).

Next, we monitored the potential for AA147 to modify KEAP1 covalently. We expressed FLAG-tagged KEAP1 (KEAP1^FT^) in HEK293 cells treated with or without AA147^alk^. We then labeled AA147^alk^ with rhodamine-azide using ‘click chemistry’ and monitored the population of rhodamine-labeled and total immunoprecipitated KEAP1^FT^ by SDS-PAGE by rhodamine fluorescence and immunoblotting, respectively. We observed dose-dependent increases in KEAP1^FT^ labeling with AA147, confirming that our compound covalently targets KEAP1 (**Fig. 5C**). Interestingly, mutating the KEAP1 sensor cysteine 151 to serine (C151S) reduced labeling, indicating that AA147 preferentially targets this cysteine to activate NRF2. Collectively, these results suggest that AA147-dependent activation of NRF2 requires metabolic activation of the AA147 2-*p*-amino cresol A-ring for covalent modification of KEAP1.

### NRF2 inhibition attenuates AA147-dependent protection against glutamate-induced toxicity

Upregulation of NRF2 transcriptional activity protects against glutamate toxicity in HT22 cells (13, 37, 41). Therefore, we asked whether the protection afforded by AA147 in this model was mediated by NRF2 activity. To test this, we co-pretreated HT22 cells with AA147 and the NRF2 inhibitor ML385 for 16 h prior to addition of glutamate. After 24 h, we then monitored cell death by Annexin V and PI staining. Co-treatment with ML385 significantly inhibited AA147-dependent reductions in glutamate-induced cell death (**Fig. 6A, Fig. S6A**). Similar results were observed by MTT assay (**Fig. 6B**). Interestingly, treatment with ML385 also blocked protection against glutamate toxicity induced by 6 h pre-treatment with AA147, indicating that protection derived from both timepoints is sensitive to pharmacologic NRF2 inhibition (**Fig. 6C**). Further, ML385 co-treatment attenuated AA147-dependent decreases in ROS in glutamate-treated HT22 cells, indicating that AA147 reductions in ROS levels can be partially attributed to NRF2 activation (**Fig. S6B**). We next shRNA-depleted *Nrf2* in HT22 cells to further define the dependence of the observed AA147-dependent protection on NRF2 activity (**Fig. S6C**). As observed with ML385, AA147 did not improve viability in glutamate-treated HT22 cells shRNA depleted of *Nrf2*, further indicating that AA147-dependent NRF2 activation is required for protection against glutamate toxicity (**Fig. 6C**). Combined, these results demonstrate that AA147 primarily mitigates glutamate-induced oxidative stress in HT22 cell through an NRF2-dependent mechanism.

**Figure 6.**
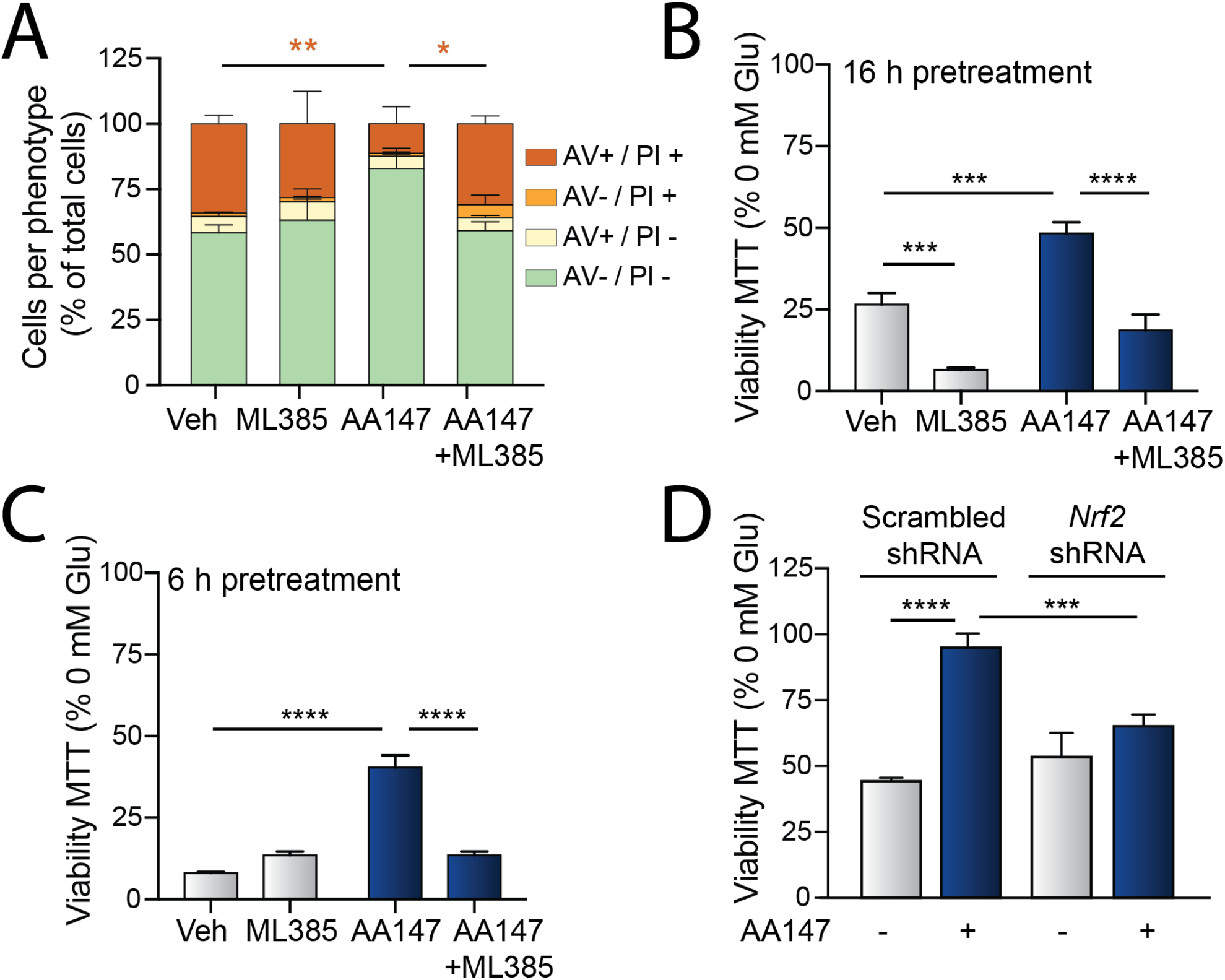
AA147 induced protection from glutamate toxicity is attenuated by NRF2 inhibition. **A**. Quantification of the percent of HT22 cells pre-treated with AA147 for 6 h and then challenged with glutamate (5 mM) for 24 h stained with Annexin V (AV) and/or propidium iodide (PI). Error bars show SEM for n=3 replicates. *p<0.05, **p<0.01 for ordinary one-way ANOVA with Tukey correction for multiple comparisons. **B**. Viability, measured by MTT, of HT22 cells pre-treated with AA147 (10 µM) for 16 h (**B**) or 6 h (**C**) in the presence or absence of the NRF2 inhibitor ML385 (5 µM) and then challenged with glutamate (5 mM) for 24 h. Viability is reported as percent relative to vehicle. ***p<0.001, ****p<0.0001 for two-way ANOVA relative to vehicle with Tukey correction for multiple comparisons. **D**. Viability, measured by MTT, of HT22 cells expressing scrambled or *Nrf2* shRNA and pre-treated for 16 h with AA147 and then challenged with glutamate (5 mM) for 24 h. Viability is reported as percent relative to vehicle. Error bars show SD for n=3. ***p<0.001, ****p<0.0001 for two-way ANOVA with Tukey correction for multiple testing.

### Concluding Remarks

Previous results showed that the proteostasis regulator compound AA147 protects against oxidative damage through the activation of the ATF6 signaling arm of the UPR (14, 15). Here, we demonstrate that AA147 protects against glutamate induced oxidative toxicity in neuronal derived HT22 cells primarily through a mechanism involving activation of the NRF2-regulated oxidative stress response. Our results indicate that AA147-dependent activation of ATF6 and NRF2 share a similar mechanism of activation involving compound oxidation to a reactive electrophile and covalent modification of protein substrates (20). However, unlike ATF6 activation, which involves AA147-dependent modification of PDIs (16, 20), AA147-dependent activation of NRF2 involves compound-dependent modification of KEAP1. This demonstrates that protective NRF2 signaling can be activated in neurons using metabolically activated compounds such as AA147. Further, our results indicate that AA147 can promote protection against oxidative insults through the activation of two distinct stress-responsive signaling pathways, the ATF6 arm of the UPR (16) and the NRF2 oxidative stress response (described herein). These results highlight the broad potential for this compound to mitigate oxidative damage in etiologically diverse diseases, including many neurodegenerative disorders.

## MATERIALS AND METHODS

### Compounds, Antibodies, and Plasmids

AA147 and associated analogs were reported previously and obtained from the Kelly Lab at Scripps Research (20). AA147 and related analogs were suspended in DMSO (20). Cells were treated with 10 µM of these compounds unless otherwise stated. PF429242 (Sigma; cat SML0667) was resuspended in water and administered at 10 µM. CP7 was obtained from the Walter Lab at UCSF, resuspended in DMSO, and administered at 5 µM. ML385 (Cayman Chemicals; cat. 21114) was resuspended in DMSO and administered at 5 µM. Glutamate stocks were prepared using glutamic acid (Acros Organics; cat. AC156211000) resuspended in water and the pH adjusted to 7.5. Equivalent water volume was used as control for all 0 mM glutamate treatments. The following antibodies were purchased and utilized in this study as indicated: NQO1 (1:1000; Abcam cat. ab80588), KDEL (1:1000; Enzo cat. ADI-SPA-827-F), tubulin (1:2000; Sigma-Aldrich T6074). The ERSE-LUC and ARE-LUC plasmids were previously described (15, 42). For viral transfection, the following plasmids were used: REV (pRSV-rev; Addgene cat. 12253), RRE (pMDL-RRE; Addgene cat. 12251),), and VSV-G (pMD2.G; Addgene cat. 12259). *ATF6* and *NRF2* shRNAs in pLKO.1 vectors were obtained from La Jolla Institute for Allergy and Immunology (LJI). The specific target sequences for viral plasmid these shRNAs are below:

*ATF6*-1-TRCN0000008447

CCGGCGAAGGGATCATCTGCTATTACTCGAGTAATAGCAGATGATCCCTTCGTTTTT

*ATF6*-TRCN0000008448

CCGGGCCATCATCATTCAGACACTACTCGAGTAGTGTCTGAATGATGATGGCTTTTT

*NRF2*-TRCN0000007555

CCGGGCTCCTACTGTGATGTGAAATCTCGAGATTTCACATCACAGTAGGAGCTTTTT

### Cell culture maintenance and shRNA depletion

HT22 cells were a kind gift from Pamela Maher at the Salk Institute. HT22 cells were split when 70% confluent and discarded after 10 passages. During experimental testing, HT22 cells were plated at a density of 5×10^3^ per well for a 96 well plate or equivalent density for larger plates. All cells were routinely tested for mycoplasma and incubated in High Glucose DMEM supplemented with 10% FBS, glutamate, and penicillin/streptomycin at 37°C and 5% CO_2_. For shRNA depletion, viruses expressing specific shRNAs were prepared as previously described (20). Briefly, 1 10 cm dish of HEK293T cells per shRNA was transiently transfected with 8 µg shRNA construct, 4 µg REV (pRSV-rev), 4 µg RRE (pMDL-RRE), 4 µg VSV-G (pMD2.G). Transfection reagents were removed after a 24 h incubation, followed by a 24 h incubation for viral production in fresh media. A 1:1 ratio of virus containing media and fresh media was added to HT22 cells for 24 h. Transfected cells were puromycin-selected (5 μg/ L) (Sigma; cat P8833) for 7 days. Knockdown was confirmed by qPCR.

### Viability Assay

Pre-treatments were administered as described and glutamate added for 24 h prior to assessing viability using Cell-Titer Glo reagent (Promega; cat. PRG7572) or MTT 3-(4,5-dimethylthiazol-2-yl)-2,5-diphenyltetrazolium bromide) (Life Sciences; cat. M6494). Cell-Titer Glo was performed following manufacturer’s protocol. MTT viability assays were performed as described elsewhere (43). Briefly, MTT was resuspended in Dulbecco’s Phosphate Buffered Saline (DPBS; Life Tech; cat. 14190235) at a concentration of 5 mg/ml and sterile filtered prior to use. Cells were treated in 100 μL of media. 10 μL of MTT solution was added to media and incubated at 37°C for 4 h. The reaction was halted by adding 100 μL of a stop solution consistenting of 10% SDS with 10 mM HCl. Cells were allowed to completely lyse by an overnight incubation at 37°C. Absorbance was measured using SPECTRAmax PLUS 384 (Molecular Devices) plate reader at OD570 with an OD630 reference.

### Propidium Iodide (PI) and Annexin V staining

PI/Annexin V staining was performed on HT22 cells treated as indicated. Cells were challenged with glutamate for 24 h. We then harvested the cells from the plate, washed with DPBS, and resuspended in 50 μL 1X Annexin V binding buffer (BD Biosciences; cat. 556454). Cells were incubated in the dark at RT with 3 μL propidium iodide (Miltenyi Biotech; cat. 130-093-233) and 3 μL FITC-Annexin V (BD Biosciences; cat. 556419) for 20 minutes and then diluted with 100 μL binding buffer. Unstained and single-channel controls were used for compensation calculations for each experiment. Flow cytometry was performed on a NovoCyte 3000 (Acea); PI was detected at ex. 488 nm, em. 615/20 nm and FITC-Annexin V using ex. 488 nm, em. 530/30 nm channel. Analysis and gating was performed using FlowJo software (BD Biosciences, San Diego).

### Quantification of ROS by DCFDA Fluorescence

Cells were plated in 24-well clear tissue culture-treated (Genesee Scientific, San Diego) plates and compounds were pre-treated as stated for 16 h. Glutamate was added for 8 h. Following glutamate incubation, cells were harvested then washed and resuspended in DPBS. CM-H2DCFDA (Invitrogen; cat. C6827) was freshly dissolved in DMSO. Cells were incubated in 5 µM CMH2DCFDA for 30 min and immediately run on a NovoCyte 3000 (Acea) using ex. 488 nm, em. 530/30 nm channel. Cytometric analysis was performed using FlowJo software (BD Biosciences, San Diego).

### Luciferase Assays

Cells were seeded at a density of 3,500 cells per well into flat white 384-well plates (Corning; cat). The following day, cells were transfected with p-TI-ARE-LUC (42) or pcDNA3.1-ERSE-LUC (15) plasmids (100 ng/well) using polyethyleneimine (PEI) at a ratio of 2:1 (PEI: DNA). Media was changed 16 h later to remove PEI. Cells were treated as indicated for 16 h then lysed by the addition of Bright-Glo (Promega). Samples were dark adapted for 20 min to stabilize signals. Luminescence was then measured in an Infinite F200 PRO plate reader (Tecan) and corrected for background signal.

### KEAP1 Modification

Cells were transiently transfected with FLAG-KEAP1 C151S or WT construct. The following day, media was changed to remove transfection agent. The following day, cells were incubated with 10 µM AA147^alk^ or otherwise as described overnight. Cells were washed with DPBS and lysed in RIPA followed by sonication. Lysate was incubated with M2-FLAG beads overnight. Beads were washed three times and FLAG-tagged proteins eluted using 3X FLAG peptide (Sigma; cat. F4799). Rhodamine-Azide labeling reactions were performed using 1.7 mM TBTA, 50 mM CuSO4, 5 mM azide, 50 mM TCEP. Protein was purified using MeOH precipitation and run on 12% polyacrylamide gel. Eluate was resuspended in laemmlil and run directly on a 12% polyacrylamide gel and immunoblotted using 1:1000 M2-FLAG antibody (Sigma; cat. F1804).

### Quantitative PCR (qPCR)

HT22 cells were treated with 10 µM AA147 or DMSO vehicle for either 6 or 16h. Cells were rinsed with PBS, lysed, and total RNA collected using the QuickRNA mini kit (Zymo) according to the manufacturer’s instructions. The relative quantification of mRNA was calculated using qPCR with reverse transcription (RT-qPCR). RNA yield was quantified using Nanodrop. cDNA was generated from 300 ng of RNA using High-Capacity cDNA Reverse Transcription Kit (Advanced Biosystems; cat. 4368814). qPCR reactions were prepared using Power SYBR Green PCR Master Mix (Applied Biosystems; cat. 4367659) and were obtained primers from Integrated DNA Technologies. Amplification reactions were run in an ABI 7900HT Fast Real Time PCR machine with an initial melting period of 95 °C for 5 min then 45 cycles of 10 s at 95 °C, 30 s at 60 °C.

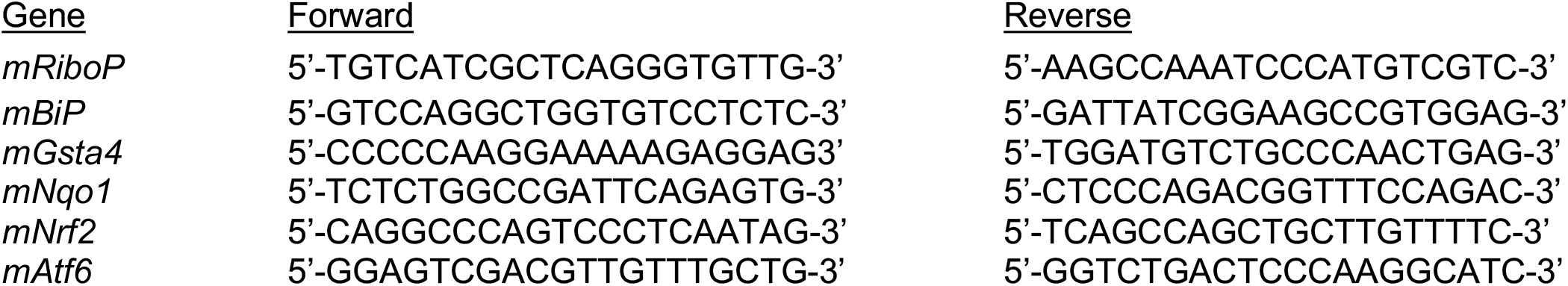

### Immunoblotting

Cell lysates were prepared as previously described (44). Briefly, cells were lysated in RIPA buffer (50 mM Tris, pH 7.5, 150 mM NaCl, 0.1% SDS, 1% Triton X-100, 0.5% deoxycholate and protease inhibitor cocktail (Roche). Total protein concentration in cellular lysates was normalized using the Bio-Rad protein assay. Lysates were then denatured with 1× Laemmli buffer + 100 mM DTT and boiled before being separated by SDS–PAGE. Samples were transferred onto nitrocellulose membranes (Bio-Rad). Membranes were then incubated overnight at 4 °C with primary antibodies diluted at 1:1000. Membranes were washed in TBST, incubated with the species appropriate IR-Dye conjugated secondary antibodies and analyzed using Odyssey Infrared Imaging System (LI-COR Biosciences). Quantification was carried out with LI-COR Image Studio software.

### RNA Sequencing

HT22 cells were treated for 16 h with 10 µM AA147 or vehicle. Cells were rinsed with DPBS, lysed and total RNA collected using the QuickRNA mini kit (Zymo) according to the manufacturer’s instructions. Transcriptional profiling using whole transcriptome RNA sequencing was conducted via BGI Americas on the BGI Proprietary platform with three biological replicates for each condition. All samples were sequenced to a minimum depth of 27 M PE 100 bp reads. Alignment of reads was performed using DNAstar Lasergene SeqManPro to the mouse genome GRCm39 assembly. Aligned reads were imported into ArrayStar 12.2 with QSeq (DNAStar Inc.) to quantify the gene expression levels. Differential expression analysis and statistical significance calculations between different conditions was assessed using DESeq2 in R, compared to vehicle treated cells. The complete RNAseq data is deposited in gene expression omnibus (GEO) as GSE178964..

### Code Availability

Code for standard open-source DESeq2 differential gene expression RNA-seq analysis used in R statistical software is available from the corresponding author upon reasonable request.

### Statistical Methods

All statistical analyses were performed using Prism 9 (GraphPad, San Diego, CA, USA) as described. The number of replicates and independent experiments for each figure panel are clearly stated in the figure legends. EC50 calculations were performed using log(agonist) vs. response variable slope four parameter nonlinear function with least squares fit.

## ACKNOWLEDGEMENTS

We thank P. Maher (Salk Institute) for generously providing HT22 cells. We thank Lara Ibrahim, Dorian Rosen, Caroline Stanton, and Gabe Kline at Scripps Research for fruitful discussions and experimental support related to the work described herein. We also thank Ryan Paxman and Jeff Kelly at Scripps Research for providing the analogs of AA147 used herein. We would also like to thank Evan Powers and Michael Petrascheck for critical reading of this manuscript. This work was supported by the NIH (AG095892 and DK107604 to RLW).

## Abbreviations

NRF2: nuclear factor erythroid 2-related factor 2
ATF6: activating transcription factor 6
UPR: unfolded protein response
ER: endoplasmic reticulum
S1P: site 1 protease
ERSE: ER-stress responsive element
ARE: antioxidant response element

## FIGURES

**Figure S1.**
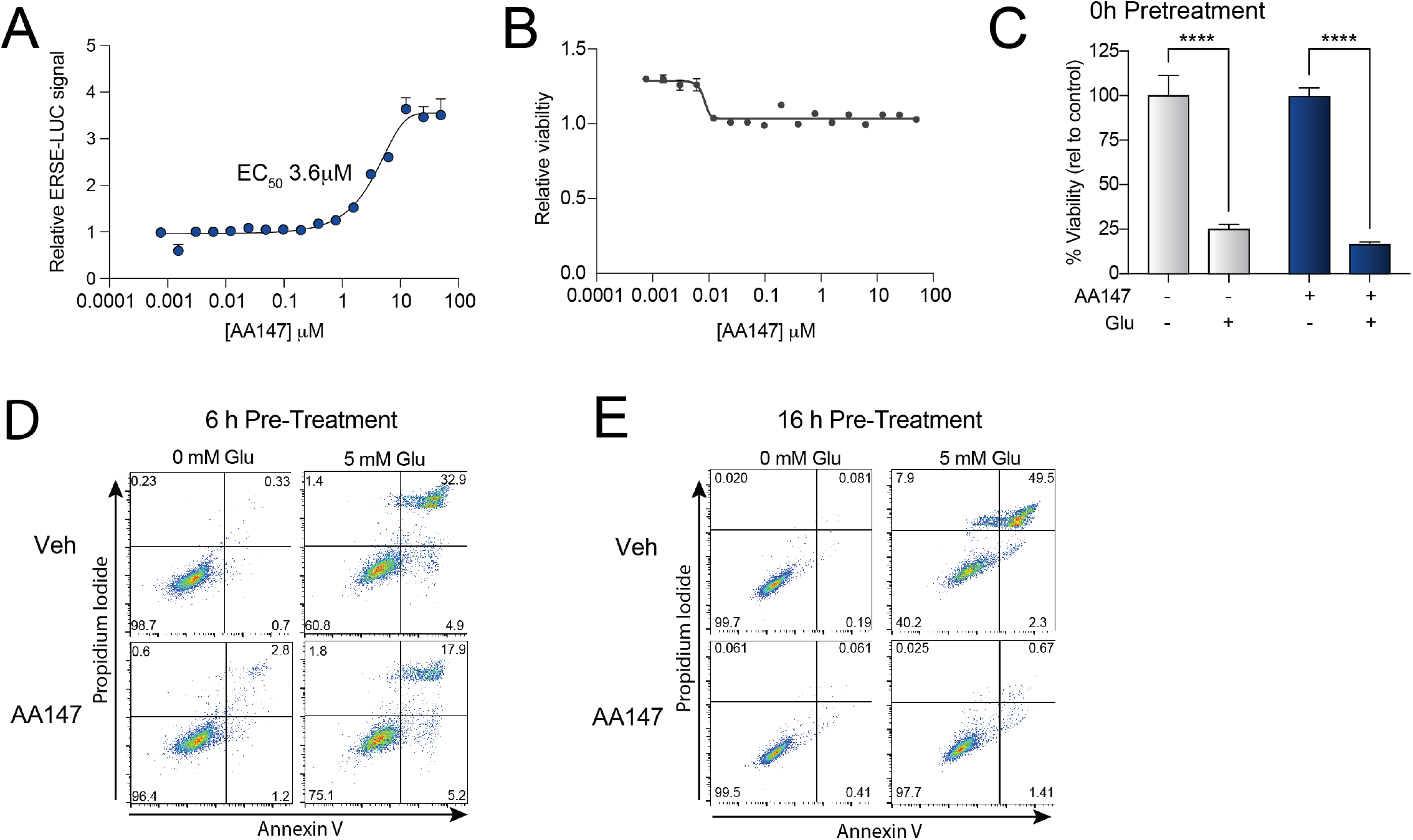
AA147 protects against glutamate-induced oxidative toxicity in HT22 cells. **A**. Luminescence in HT22 cells transiently expressing the ERSE-LUC reporter treated with the indicated concentration of AA147 for 16 h. Luminescence is shown as fold-change relative to vehicle. Error bars show SEM for n=20 replicates across two independent experiments. **B**. CellTiterGlo luminescence in HT22 cells treated with the indicated concentration of AA147 for 16 h. Luminescence is shown as a fold-change relative to vehicle. Error bars show SEM for n=20 replicates across two independent experiments. **C**. Viability, measured by MTT, of HT22 cells treated concurrently with AA147 (10 µM) and glutamate (5 mM) for 24 h. Viability is shown as percent relative to vehicle treated cells with no glutamate added. Error bars show SEM for n=3. ****p<0.001 for two-way ANOVA with Tukey correction for multiple testing. **D,E**. Representative flow cytometry plots, with single-channel compensation correction, showing Annexin V and propidium iodide staining of HT22 cells pre-treated with AA147 (10 µM) for 6 h (**D**) or 16 h (**E**) and then challenged with glutamate (5 mM) for 24 h.

**Figure S2.**
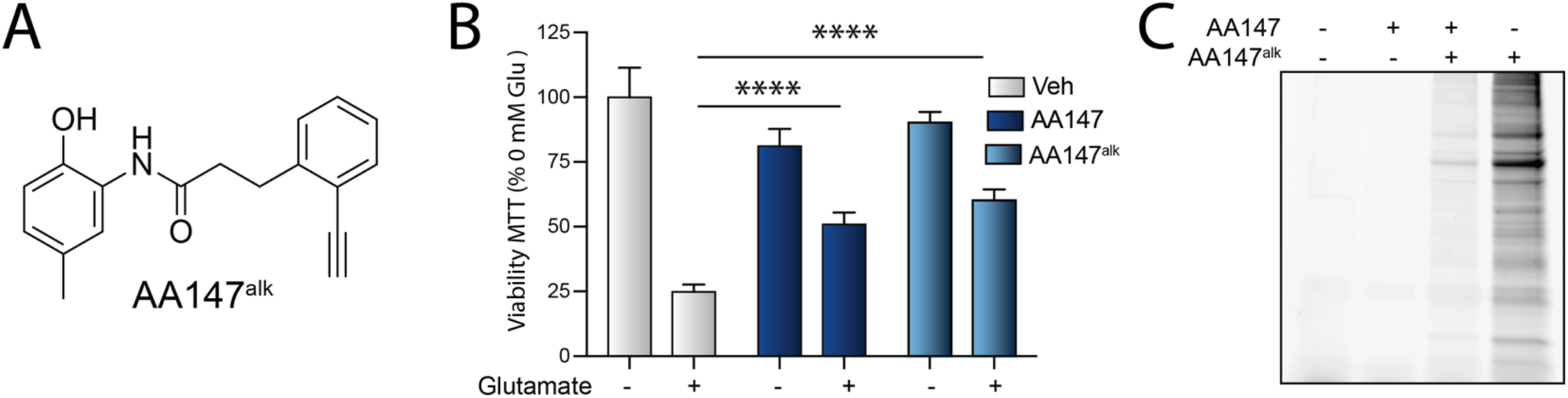
The 2-*p*-amino cresol substructure of AA147 is required to protect HT22 cells against glutamate induced oxidative toxicity. **A**. Structure of AA147^alk^. **B**. Viability of HT22 cells, measured by MTT, pre-treated with AA147 (10 µM) or AA147^alk^ (10 µM) for 16 h and then challenged with glutamate (5 mM) for 24 h. Viability is reported as percent viability normalized to vehicle treated cells in the absence of glutamate. Error bars show SD for n=3 replicates. ****p<0.0001 for ordinary one-way ANOVA against vehicle control with Dunnett correction for multiple comparisons. **C**. SDS-PAGE of lysates prepared from HT22 cells treated for 16 h with AA147 (50 µM) and/or AA147^alk^ (10 µM). Proteins modified with AA147^alk^ were conjugated to a rhodamine dye using ‘click’ chemistry and imaged by rhodamine fluorescence.

**Figure S3.**
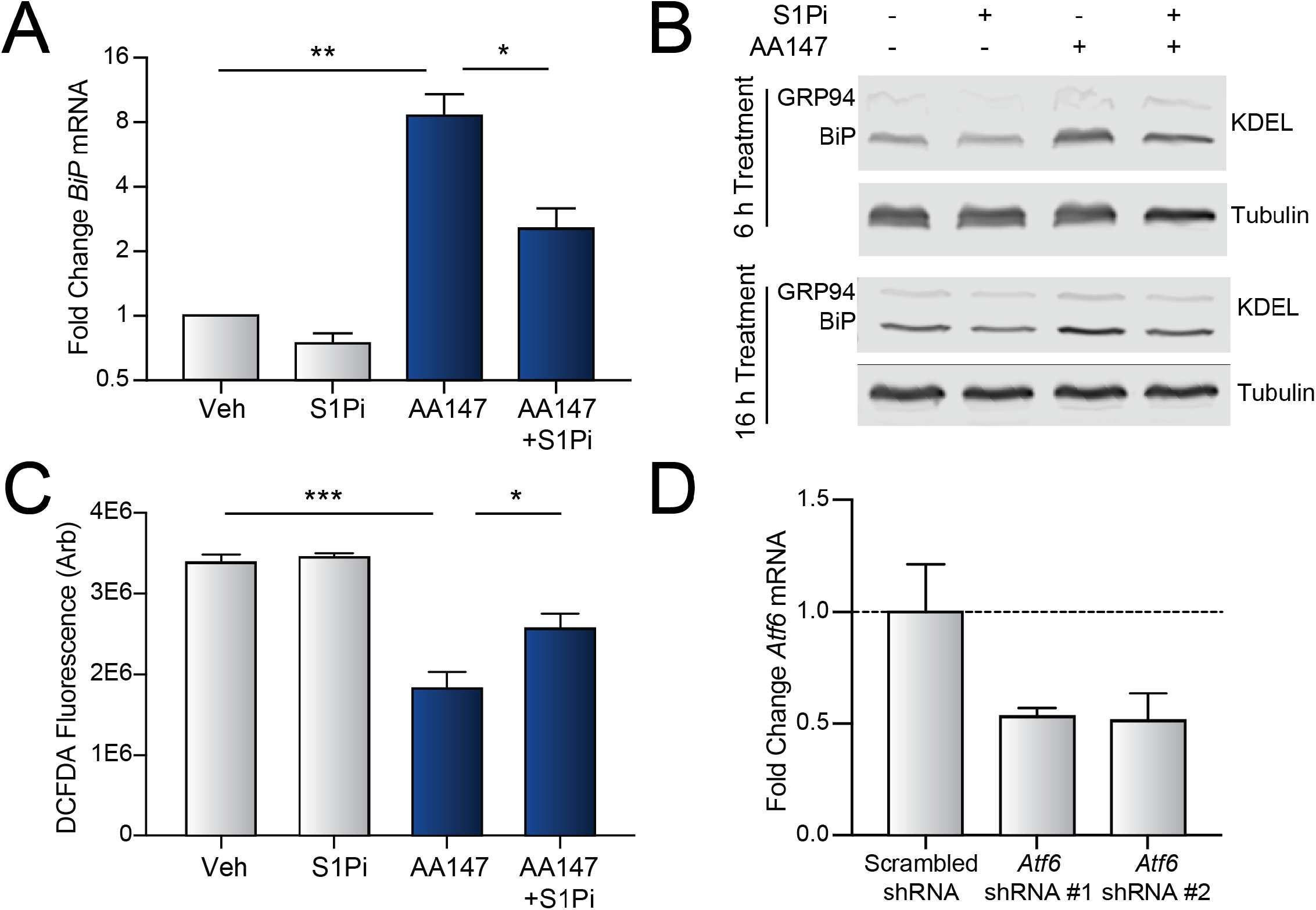
AA147-dependent activation of ATF6 modestly contributes to the AA147-dependent protection of HT22 cells against glutamate induced oxidative toxicity. **A**. Expression of the ATF6 target gene *BiP*/*Hspa5*, measured by qPCR, in HT22 cells treated with vehicle or AA147 (10 µM) in the presence or absence of S1Pi (10 µM) for 6 h. Error bars show SEM for n=3. *p<0.05, **p<0.01 for two-way ANOVA with Tukey correction for multiple testing. **B**. Representative immunoblots showing BiP and GRP94 (both detected by the KDEL antibody) in lysates prepared from HT22 cells pre-treated with vehicle or AA147 (10 µM) for 6 h and then incubated for an additional 12 h (top) or 18 h (bottom). Tubulin is shown as a loading control. **C**. Mean CM-H2DCFDA fluorescence of HT22 cells pre-treated with AA147 (10 µM) and/or S1Pi (10 µM) for 16 h and then challenged with glutamate (5 mM) for 8 h, as indicated. Error bars show SEM for n=3 replicates. *p<0.05, ***p<0.005 for two-way ANOVA with Tukey correction for multiple testing. **D**. Expression of *Atf6*, measured by qPCR, in HT22 cells stably expressing scrambled or *Atf6* shRNA. Error bars show +/- 95% confidence intervals for n=3 technical replicates.

**Figure S4.**
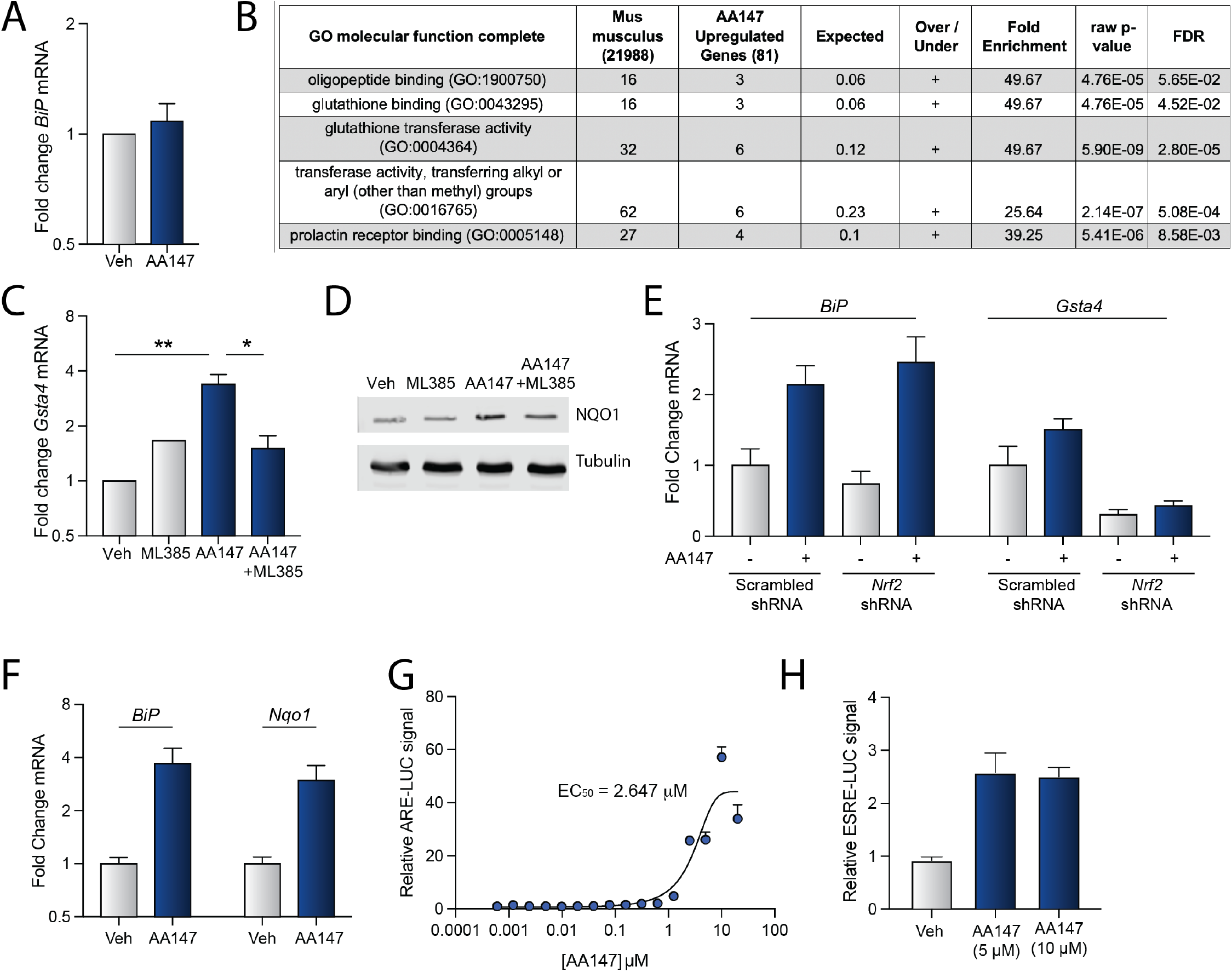
AA147 induces NRF2-dependent upregulation of oxidative stress response genes in HT22 cells. **A**. Expression of the ATF6 target gene *BiP*/*Hspa5*, measured by qPCR, in HT22 cells treated with vehicle or AA147 (10 µM) for 16 h. Error bars show SEM for n=3. **B**. Gene ontology (GO) for genes shown to be increased with a fold change >2 and adj p <0.01 in AA147-treated HT22 cells measured by RNA-seq. **C**. Expression of the NRF2 target gene *Gsta4* in HT22 cells treated with vehicle or AA147 (10 µM) in the presence or absence of ML385 (5 µM) for 16 h. Error bars show SEM for n=3. *p<0.05, **p<0.01 for two-way ANOVA with Tukey correction for multiple testing. **D**. Immunoblot of NQO1 in lysates prepared from HT22 cells treated with vehicle or AA147 (10 µM) for 18 h. Tubulin is shown as a loading control. **E**. Expression of the ATF6 target gene *BiP* and the NRF2 target gene *Gsta4* in HT22 cells stably transfected with scrambled or *Nrf2* shRNA and treated for 16 h with vehicle or AA147 (10 µM). Error bars show +/- 95% confidence intervals for n=3 technical replicates. **F**. Expression of the ATF6 target gene *Bip/Hspa5* and the NRF2 target gene *Nqo1*, measured by qPCR, in DIV27 primary mouse cortical neurons treated with vehicle or AA147 (10 µM) for 6 h. Error bars show +/- 95% confidence intervals for n=3 technical replicates. **G**. Luminescence in IMR32 cells transiently expressing the ARE-LUC NRF2 reporter treated with the indicated concentration of AA147 for 16 h. Luminescence is shown as percent of vehicle. Error bars show SEM for n=3. **H**. Luminescence in IMR32 cells transiently expressing the ERSE-LUC ATF6 reporter treated with the indicated concentration of AA147 for 16 h. Luminescence is shown as percent of vehicle. Error bars show SEM for n=3.

**Figure S5.**
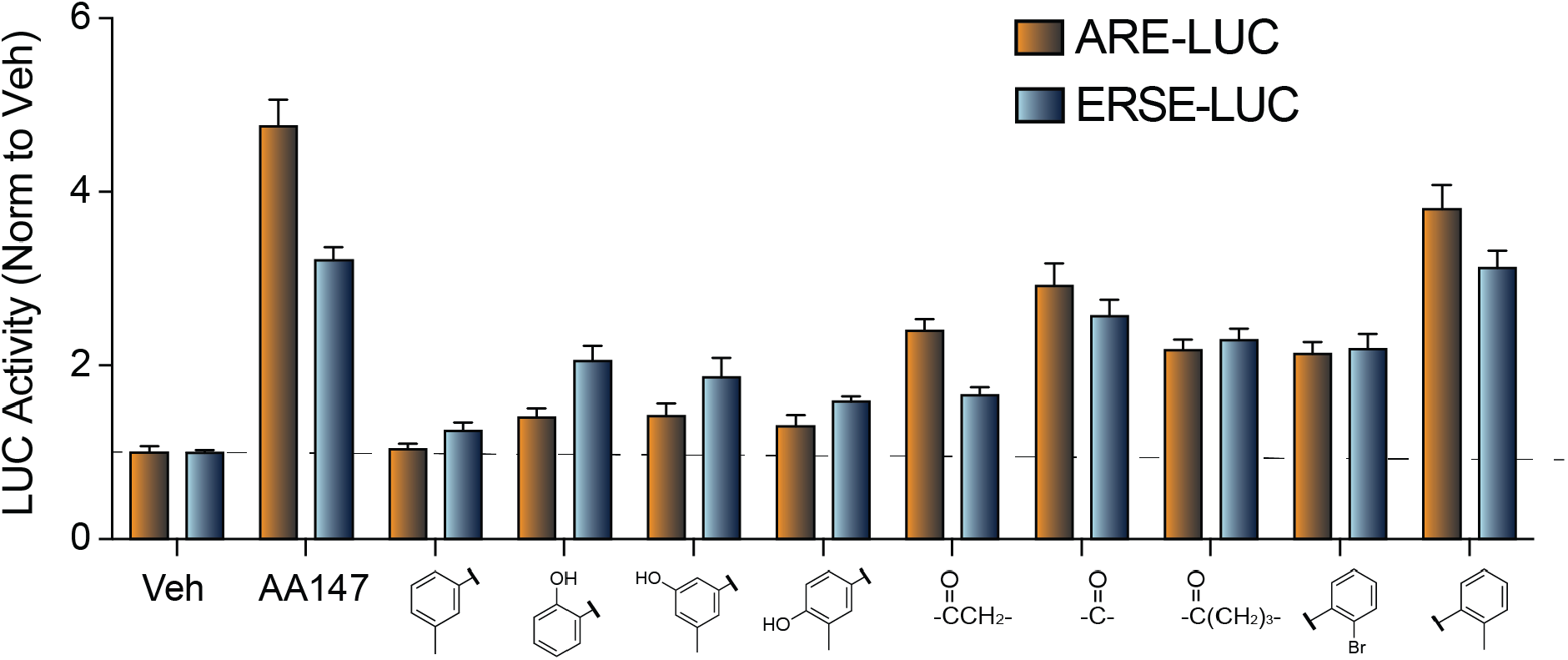
AA147 covalently modifies KEAP1, a regulator of NRF2 transcriptional activity. Comparison between activation of the ATF6-responsive ERSE-LUC reporter and the NRF2-responsive ARE-LUC reporter in HT22 cells expressing the indicated reporter treated with the different AA147 analog (10 µM) for 16 h. Luminescence is reported as fold change relative to vehicle. Error bars show SEM for n=20 replicates across two independent experiments.

**Figure S6.**
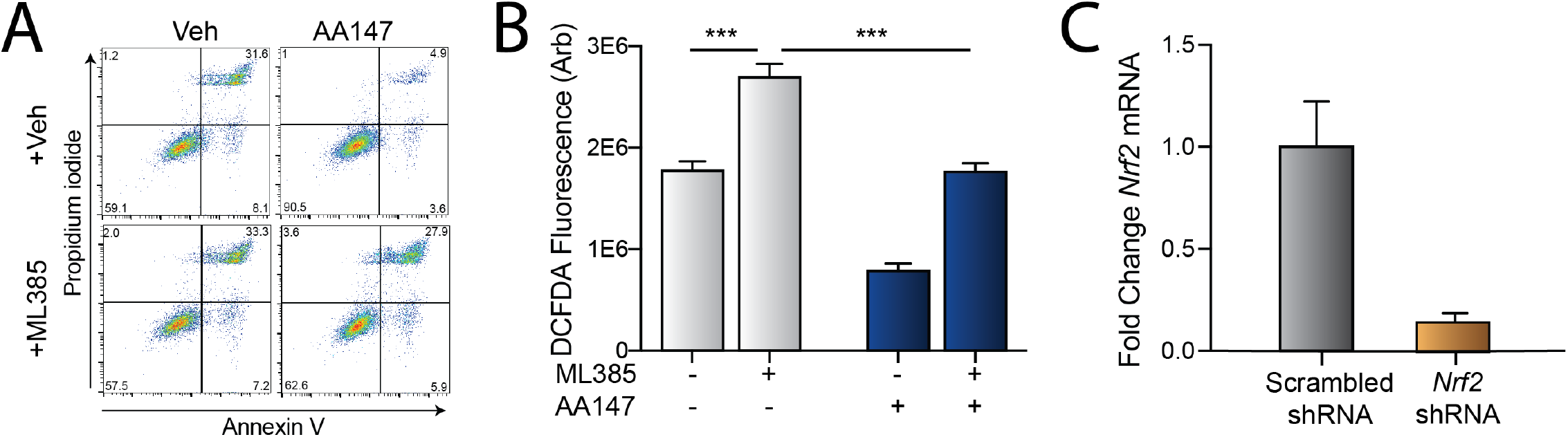
AA147 induced protection from glutamate toxicity is attenuated by NRF2 inhibition. **A**. Representative propidium iodide and Annexin V staining of HT22 cells pre-treated with AA147 (10 µM) for 6 h in the presence or absence of ML384 (5 µM) and then challenged with glutamate (5 mM) for 24 h. **B**. Mean CM-H2DCFDA fluorescence of HT22 cells pre-treated with AA147 (10 µM) and/or ML385 (10 µM) for 6 h and then challenged with glutamate (5 mM) for 8 h, as indicated. Error bars show SEM for n=3 replicates. ***p<0.001 for two-way ANOVA with Tukey correction for multiple testing. **C**. Expression of *Nrf2*, measured by qPCR, in HT22 cells expressing scrambled or *Nrf2* shRNA. Error bars show +/- 95% confidence intervals for n=3 technical replicates.

## SUPPLEMENTARY TABLE LEGENDS (see included Excel sheets for the Tables)

**Supplementary Table S1. RNAseq of HT22 cells treated with AA147 for 16 h**. The included excel spread shows the DESEQ2 analysis of RNAseq of HT22 cells treated with vehicle or AA147 (10 µM) for 16 h. The complete RNAseq data is deposited in gene expression omnibus (GEO) as GSE178964..

**Supplementary Table S2. Geneset-based stress-signaling pathway profiling of HT22 cells treated with AA147 for 16 h**. Excel spreadsheet showing the DESEQ2 analysis of select genes identified to be activated downstream of stress-responsive signaling pathways including the ATF6, IRE1/XBP1s, and PERK signaling arms of the unflded protein response (UPR), the HSF1-regulated heat shock response (HSR), the oxidative stress response (OSR), the hypoxia stress response, the NFkB inflammatory stress response, and the mitochondrial unfolded protein response (UPR^mt^), as well as a set of control genes, from RNAseq of HT22 cells treated with AA147 (10 µM, 16 h; see **Supplemental Table S1**). Genesets used in this analysis were defined in Grandjean et al (2019) *ACS Chem Biol*.

## REFERENCES

1. Bano D, Young KW, Guerin CJ, Lefeuvre R, Rothwell NJ, Naldini L, Rizzuto R, Carafoli E, Nicotera P. Cleavage of the plasma membrane Na+/Ca2+ exchanger in excitotoxicity. Cell. 2005;120(2):275–85. Epub 2005/02/01. doi: 10.1016/j.cell.2004.11.049. PubMed PMID: 15680332.

2. Armada-Moreira A, Gomes JI, Pina CC, Savchak OK, Gonçalves-Ribeiro J, Rei N, Pinto S, Morais TP, Martins RS, Ribeiro FF, Sebastião AM, Crunelli V, Vaz SH. Going the Extra (Synaptic) Mile: Excitotoxicity as the Road Toward Neurodegenerative Diseases. Frontiers in cellular neuroscience. 2020;14(90). doi: 10.3389/fncel.2020.00090.

3. Dirnagl U, Iadecola C, Moskowitz MA. Pathobiology of ischaemic stroke: an integrated view. Trends in Neurosciences. 1999;22(9):391–7. doi: https://doi.org/10.1016/S0166-2236(99)01401-0.

4. Ankarcrona M, Dypbukt JM, Bonfoco E, Zhivotovsky B, Orrenius S, Lipton SA, Nicotera P. Glutamate-induced neuronal death: a succession of necrosis or apoptosis depending on mitochondrial function. Neuron. 1995;15(4):961–73. Epub 1995/10/01. doi: 10.1016/0896-6273(95)90186-8. PubMed PMID: 7576644.

5. Kritis AA, Stamoula EG, Paniskaki KA, Vavilis TD. Researching glutamate - induced cytotoxicity in different cell lines: a comparative/collective analysis/study. Frontiers in cellular neuroscience. 2015;9:91. Epub 2015/04/09. doi: 10.3389/fncel.2015.00091. PubMed PMID: 25852482; PMCID: PMC4362409.

6. Maher P, Davis JB. The role of monoamine metabolism in oxidative glutamate toxicity. The Journal of neuroscience : the official journal of the Society for Neuroscience. 1996;16(20):6394–401. Epub 1996/10/15. PubMed PMID: 8815918.

7. Soria FN, Perez-Samartin A, Martin A, Gona KB, Llop J, Szczupak B, Chara JC, Matute C, Domercq M. Extrasynaptic glutamate release through cystine/glutamate antiporter contributes to ischemic damage. The Journal of clinical investigation. 2014;124(8):3645–55. Epub 2014/07/19. doi: 10.1172/jci71886. PubMed PMID: 25036707; PMCID: PMC4109556.

8. Davis JB, Maher P. Protein kinase C activation inhibits glutamate-induced cytotoxicity in a neuronal cell line. Brain research. 1994;652(1):169–73. Epub 1994/07/25. PubMed PMID: 7953717.

9. Long JM, Holtzman DM. Alzheimer Disease: An Update on Pathobiology and Treatment Strategies. Cell. 2019;179(2):312–39. Epub 2019/10/01. doi: 10.1016/j.cell.2019.09.001. PubMed PMID: 31564456; PMCID: PMC6778042.

10. Berthier ML, Green C, Lara JP, Higueras C, Barbancho MA, Dávila G, Pulvermüller F. Memantine and constraint-induced aphasia therapy in chronic poststroke aphasia. Annals of Neurology. 2009;65(5):577–85. doi: https://doi.org/10.1002/ana.21597.

11. Trotman M, Vermehren P, Gibson CL, Fern R. The dichotomy of memantine treatment for ischemic stroke: dose-dependent protective and detrimental effects. Journal of cerebral blood flow and metabolism : official journal of the International Society of Cerebral Blood Flow and Metabolism. 2015;35(2):230–9. Epub 2014/11/20. doi: 10.1038/jcbfm.2014.188. PubMed PMID: 25407270; PMCID: PMC4426739.

12. Maher P, Currais A, Schubert D. Using the Oxytosis/Ferroptosis Pathway to Understand and Treat Age-Associated Neurodegenerative Diseases. Cell chemical biology. 2020;27(12):1456–71. Epub 2020/11/12. doi: 10.1016/j.chembiol.2020.10.010. PubMed PMID: 33176157; PMCID: PMC7749085.

13. Lewerenz J, Maher P. Control of redox state and redox signaling by neural antioxidant systems. Antioxid Redox Signal. 2011;14(8):1449–65. Epub 2010/09/04. doi: 10.1089/ars.2010.3600. PubMed PMID: 20812872.

14. Blackwood EA, Azizi K, Thuerauf DJ, Paxman RJ, Plate L, Kelly JW, Wiseman RL, Glembotski CC. Pharmacologic ATF6 activation confers global protection in widespread disease models by reprograming cellular proteostasis. Nature communications. 2019;10(1):187. Epub 2019/01/16. doi: 10.1038/s41467-018-08129-2. PubMed PMID: 30643122; PMCID: PMC6331617.

15. Plate L, Cooley CB, Chen JJ, Paxman RJ, Gallagher CM, Madoux F, Genereux JC, Dobbs W, Garza D, Spicer TP, Scampavia L, Brown SJ, Rosen H, Powers ET, Walter P, Hodder P, Wiseman RL, Kelly JW. Small molecule proteostasis regulators that reprogram the ER to reduce extracellular protein aggregation. eLife. 2016;5. Epub 2016/07/21. doi: 10.7554/eLife.15550. PubMed PMID: 27435961; PMCID: PMC4954754.

16. Glembotski CC, Rosarda JD, Wiseman RL. Proteostasis and Beyond: ATF6 in Ischemic Disease. Trends in Molecular Medicine. 2019;25(6):538–50. doi: https://doi.org/10.1016/j.molmed.2019.03.005.

17. Ye J, Rawson RB, Komuro R, Chen X, Davé UP, Prywes R, Brown MS, Goldstein JL. ER Stress Induces Cleavage of Membrane-Bound ATF6 by the Same Proteases that Process SREBPs. Molecular Cell. 2000;6(6):1355–64. doi: https://doi.org/10.1016/S1097-2765(00)00133-7.

18. Shoulders MD, Ryno LM, Genereux JC, Moresco JJ, Tu PG, Wu C, Yates JR, 3rd, Su AI, Kelly JW, Wiseman RL. Stress-independent activation of XBP1s and/or ATF6 reveals three functionally diverse ER proteostasis environments. Cell reports. 2013;3(4):1279–92. Epub 2013/04/16. doi: 10.1016/j.celrep.2013.03.024. PubMed PMID: 23583182; PMCID: PMC3754422.

19. Adachi Y, Yamamoto K, Okada T, Yoshida H, Harada A, Mori K. ATF6 is a transcription factor specializing in the regulation of quality control proteins in the endoplasmic reticulum. Cell Struct Funct. 2008;33(1):75–89. Epub 2008/03/25. doi: 10.1247/csf.07044. PubMed PMID: 18360008.

20. Paxman R, Plate L, Blackwood EA, Glembotski C, Powers ET, Wiseman RL, Kelly JW. Pharmacologic ATF6 activating compounds are metabolically activated to selectively modify endoplasmic reticulum proteins. eLife. 2018;7. Epub 2018/08/08. doi: 10.7554/eLife.37168. PubMed PMID: 30084354; PMCID: PMC6080950.

21. Wang S, Kaufman RJ. The impact of the unfolded protein response on human disease. The Journal of cell biology. 2012;197(7):857–67. Epub 2012/06/27. doi: 10.1083/jcb.201110131. PubMed PMID: 22733998; PMCID: PMC3384412.

22. Martindale JJ, Fernandez R, Thuerauf D, Whittaker R, Gude N, Sussman MA, Glembotski CC. Endoplasmic Reticulum Stress Gene Induction and Protection From Ischemia/Reperfusion Injury in the Hearts of Transgenic Mice With a Tamoxifen-Regulated Form of ATF6. Circulation Research. 2006;98(9):1186–93. doi: doi:10.1161/01.RES.0000220643.65941.8d.

23. Chen John J, Genereux Joseph C, Qu S, Hulleman John D, Shoulders Matthew D, Wiseman RL. ATF6 Activation Reduces the Secretion and Extracellular Aggregation of Destabilized Variants of an Amyloidogenic Protein. Chemistry & Biology. 2014;21(11):1564–74. doi: https://doi.org/10.1016/j.chembiol.2014.09.009.

24. Kroeger H, Grimsey N, Paxman R, Chiang WC, Plate L, Jones Y, Shaw PX, Trejo J, Tsang SH, Powers E, Kelly JW, Wiseman RL, Lin JH. The unfolded protein response regulator ATF6 promotes mesodermal differentiation. Science signaling. 2018;11(517). Epub 2018/02/15. doi: 10.1126/scisignal.aan5785. PubMed PMID: 29440509; PMCID: PMC5957084.

25. Shen Y, Li R, Yu S, Zhao Q, Wang Z, Sheng H, Yang W. Activation of the ATF6 (Activating Transcription Factor 6) Signaling Pathway in Neurons Improves Outcome After Cardiac Arrest in Mice. Journal of the American Heart Association. 2021;10(12):e020216. Epub 2021/06/12. doi: 10.1161/jaha.120.020216. PubMed PMID: 34111943.

26. Rius B, Mesgarzadeh JS, Romine IC, Paxman RJ, Kelly JW, Wiseman RL. Pharmacologic targeting of plasma cell endoplasmic reticulum proteostasis to reduce amyloidogenic light chain secretion. Blood Advances. 2021;5(4):1037–49. doi: 10.1182/bloodadvances.2020002813.

27. Almasy KM, Davies JP, Lisy SM, Tirgar R, Tran SC, Plate L. Small-molecule endoplasmic reticulum proteostasis regulator acts as a broad-spectrum inhibitor of dengue and Zika virus infections. Proceedings of the National Academy of Sciences. 2021;118(3):e2012209118. doi: 10.1073/pnas.2012209118.

28. Fukui M, Song JH, Choi J, Choi HJ, Zhu BT. Mechanism of glutamate-induced neurotoxicity in HT22 mouse hippocampal cells. European journal of pharmacology. 2009;617(1-3):1–11. Epub 2009/07/08. doi: 10.1016/j.ejphar.2009.06.059. PubMed PMID: 19580806.

29. Hawkins JL, Robbins MD, Warren LC, Xia D, Petras SF, Valentine JJ, Varghese AH, Wang IK, Subashi TA, Shelly LD, Hay BA, Landschulz KT, Geoghegan KF, Harwood HJ, Jr. Pharmacologic inhibition of site 1 protease activity inhibits sterol regulatory element-binding protein processing and reduces lipogenic enzyme gene expression and lipid synthesis in cultured cells and experimental animals. The Journal of pharmacology and experimental therapeutics. 2008;326(3):801–8. Epub 2008/06/26. doi: 10.1124/jpet.108.139626. PubMed PMID: 18577702.

30. Chanas SA, Jiang Q, McMahon M, McWalter GK, McLellan LI, Elcombe CR, Henderson CJ, Wolf CR, Moffat GJ, Itoh K, Yamamoto M, Hayes JD. Loss of the Nrf2 transcription factor causes a marked reduction in constitutive and inducible expression of the glutathione S-transferase Gsta1, Gsta2, Gstm1, Gstm2, Gstm3 and Gstm4 genes in the livers of male and female mice. The Biochemical journal. 2002;365(Pt 2):405–16. Epub 2002/05/07. doi: 10.1042/bj20020320. PubMed PMID: 11991805; PMCID: PMC1222698.

31. Chen QM, Maltagliati AJ. Nrf2 at the heart of oxidative stress and cardiac protection. Physiological genomics. 2018;50(2):77–97. Epub 2017/12/01. doi: 10.1152/physiolgenomics.00041.2017. PubMed PMID: 29187515; PMCID: PMC5867612.

32. Hayes JD, Dinkova-Kostova AT. The Nrf2 regulatory network provides an interface between redox and intermediary metabolism. Trends in Biochemical Sciences. 2014;39(4):199–218. doi: https://doi.org/10.1016/j.tibs.2014.02.002.

33. Grandjean JMD, Plate L, Morimoto RI, Bollong MJ, Powers ET, Wiseman RL. Deconvoluting Stress-Responsive Proteostasis Signaling Pathways for Pharmacologic Activation Using Targeted RNA Sequencing. ACS chemical biology. 2019;14(4):784–95. Epub 2019/03/02. doi: 10.1021/acschembio.9b00134. PubMed PMID: 30821953; PMCID: PMC6474822.

34. Ibrahim L, Mesgarzadeh J, Xu I, Powers ET, Wiseman RL, Bollong MJ. Defining the Functional Targets of Cap’n’collar Transcription Factors NRF1, NRF2, and NRF3. Antioxidants. 2020;9(10):1025. PubMed PMID: doi:10.3390/antiox9101025.

35. Itoh K, Chiba T, Takahashi S, Ishii T, Igarashi K, Katoh Y, Oyake T, Hayashi N, Satoh K, Hatayama I, Yamamoto M, Nabeshima Y-i. An Nrf2/Small Maf Heterodimer Mediates the Induction of Phase II Detoxifying Enzyme Genes through Antioxidant Response Elements. Biochemical and biophysical research communications. 1997;236(2):313–22. doi: https://doi.org/10.1006/bbrc.1997.6943.

36. Brandes MS, Gray NE. NRF2 as a Therapeutic Target in Neurodegenerative Diseases. ASN Neuro. 2020;12:1759091419899782. doi: 10.1177/1759091419899782. PubMed PMID: 31964153.

37. Lewerenz J, Albrecht P, Tien ML, Henke N, Karumbayaram S, Kornblum HI, Wiedau-Pazos M, Schubert D, Maher P, Methner A. Induction of Nrf2 and xCT are involved in the action of the neuroprotective antibiotic ceftriaxone in vitro. J Neurochem. 2009;111(2):332–43. Epub 2009/08/22. doi: 10.1111/j.1471-4159.2009.06347.x. PubMed PMID: 19694903.

38. Singh A, Venkannagari S, Oh KH, Zhang YQ, Rohde JM, Liu L, Nimmagadda S, Sudini K, Brimacombe KR, Gajghate S, Ma J, Wang A, Xu X, Shahane SA, Xia M, Woo J, Mensah GA, Wang Z, Ferrer M, Gabrielson E, Li Z, Rastinejad F, Shen M, Boxer MB, Biswal S. Small Molecule Inhibitor of NRF2 Selectively Intervenes Therapeutic Resistance in KEAP1-Deficient NSCLC Tumors. ACS chemical biology. 2016;11(11):3214–25. Epub 2016/10/18. doi: 10.1021/acschembio.6b00651. PubMed PMID: 27552339; PMCID: PMC5367156.

39. Canning P, Sorrell FJ, Bullock AN. Structural basis of Keap1 interactions with Nrf2. Free radical biology & medicine. 2015;88(Pt B):101–7. Epub 2015/06/10. doi: 10.1016/j.freeradbiomed.2015.05.034. PubMed PMID: 26057936; PMCID: PMC4668279.

40. Zhang DD, Hannink M. Distinct cysteine residues in Keap1 are required for Keap1-dependent ubiquitination of Nrf2 and for stabilization of Nrf2 by chemopreventive agents and oxidative stress. Molecular and cellular biology. 2003;23(22):8137–51. Epub 2003/10/31. doi: 10.1128/mcb.23.22.8137-8151.2003. PubMed PMID: 14585973; PMCID: PMC262403.

41. Fischer W, Currais A, Liang Z, Pinto A, Maher P. Old age-associated phenotypic screening for Alzheimer’s disease drug candidates identifies sterubin as a potent neuroprotective compound from Yerba santa. Redox Biology. 2019;21:101089. doi: https://doi.org/10.1016/j.redox.2018.101089.

42. Bollong MJ, Lee G, Coukos JS, Yun H, Zambaldo C, Chang JW, Chin EN, Ahmad I, Chatterjee AK, Lairson LL, Schultz PG, Moellering RE. A metabolite-derived protein modification integrates glycolysis with KEAP1-NRF2 signalling. Nature. 2018;562(7728):600–4. Epub 2018/10/17. doi: 10.1038/s41586-018-0622-0. PubMed PMID: 30323285.

43. Riss TL MR, Niles AL, et al.. Cell Viability Assays. In: Markossian S SG, Grossman A, et al., editor. Assay Guidance Manual Bethesda (MD): Eli Lilly & Company and the National Center for Advancing Translational Sciences; 2013 May 1 [Updated 2016 Jul 1].

44. Grandjean JMD, Madhavan A, Cech L, Seguinot BO, Paxman RJ, Smith E, Scampavia L, Powers ET, Cooley CB, Plate L, Spicer TP, Kelly JW, Wiseman RL. Pharmacologic IRE1/XBP1s activation confers targeted ER proteostasis reprogramming. Nature chemical biology. 2020. Epub 2020/07/22. doi: 10.1038/s41589-020-0584-z. PubMed PMID: 32690944.

